# Cross-Species Biomechanical Determinants of Shape Diversity

**DOI:** 10.1101/2025.02.07.637025

**Authors:** Richard Bailleul, Nicolas Cuny, Diana Khoromoskaia, Soham Basu, Giulia Bergamini, Paolo Cucurachi, Sebastian Rupp, Annika Guse, Camille Curantz, Natalie Swinhoe, Phillip Cleves, Jamie Craggs, Sosuke Fujita, Yu-ichiro Nakajima, Petrus J. Steenbergen, Alba Diz-Muñoz, Guillaume Salbreux, Aissam Ikmi

**Author notes:** Corresponding Authors: Guillaume Salbreux and Aissam Ikmi. These authors contributed equally to this work (Richard Bailleul, Nicolas Cuny).

## Abstract

How complex molecular mechanisms translate into diverse multicellular shapes remains unclear. By leveraging the bi-layered architecture of six cnidarian species that diverged 500 million years ago, we show that modularity in supracellular mechanics governs larval shape diversity. Using active surface theory, quantitative imaging, and an inducible genetic system, we identify species-specific variations in three biomechanical modules. Basally aligned stress fibers drive axial elongation, while oral geometry and aboral rigidity define shape polarity. Remarkably, manipulating these modules transforms one species’ shape into another, demonstrating the causal relationship between module variation and shape diversity. Our analysis also uncovers instances of mechanical redundancies, where distinct module combinations generate similar shapes. These findings provide a general framework for how molecular complexity funnels into mesoscale mechanical determinants shaping morphological diversity.

## INTRODUCTION

The study of morphological evolution has traditionally emphasized the genetic underpinnings of diversity (*1, 2*). However, the development of form results from a complex interplay between genetic patterning, the spatial and temporal variation of gene expression defining cell identity, and morphogenesis, the physical process that dynamically shapes tissues through collective cellular activities (*3, 4*). While alterations in both patterning and post-patterning processes have been linked to morphological variation, most studies have focused on broad comparative molecular profiling, discrete molecular changes (*5*–*9*), or isolated cellular events such as cell shape changes (*10*), proliferation (*11*), and apoptosis (*12*). Yet, a significant gap remains in understanding how genetic diversity translates into biomechanical processes that sculpt species-specific shapes, and whether general biophysical principles govern shape evolution across species.

Cnidarians—including sea anemones, corals, and jellyfish—provide an ideal system to tackle this challenge (*13*). Their relatively simple body plan is characterized by diploblastic tissue layers, reducing the complexity seen in triploblastic bilaterians. Most cnidarians have a free-swimming, ciliated larval stage called the planula, which is critical for dispersal and habit colonization. These axisymmetric planulae exhibit species-specific variations in shape and other features, such as the presence or absence of an oral opening and pharynx (*14, 15*). By leveraging the diversity and morphological simplicity of planula larvae, we combine theory and experiments to explore how genetic regulation, tissue mechanics, and morphogenesis interact to drive shape diversity.

## RESULTS

### Shape diversity of planulae

To quantify morphological changes during the embryo-larva transition in cnidarians, we conducted a comparative shape analysis of six species (Fig. 1A), each exhibiting distinct planula morphologies (Fig. 1B and fig. S1A). In the Anthozoa class, our study included two sea anemones from the Actinaria order - *Nematostella vectensis* (*16*) - and *Exaiptasia pallida* (commonly known as *Aiptasia*) (*17*), as well as two reef-building hard corals from the Scleractinia order—*Acropora millepora* (*18*) and *Galaxea fascicularis* (*19*). Within the Medusozoa class, we examined two representatives of the Anthoathecata order: the colonial hydroid *Hydractinia symbiolongicarpus* (*20*) and *Cladonema pacificum*, which exhibits a medusa stage (*21*).

**Fig. 1.**
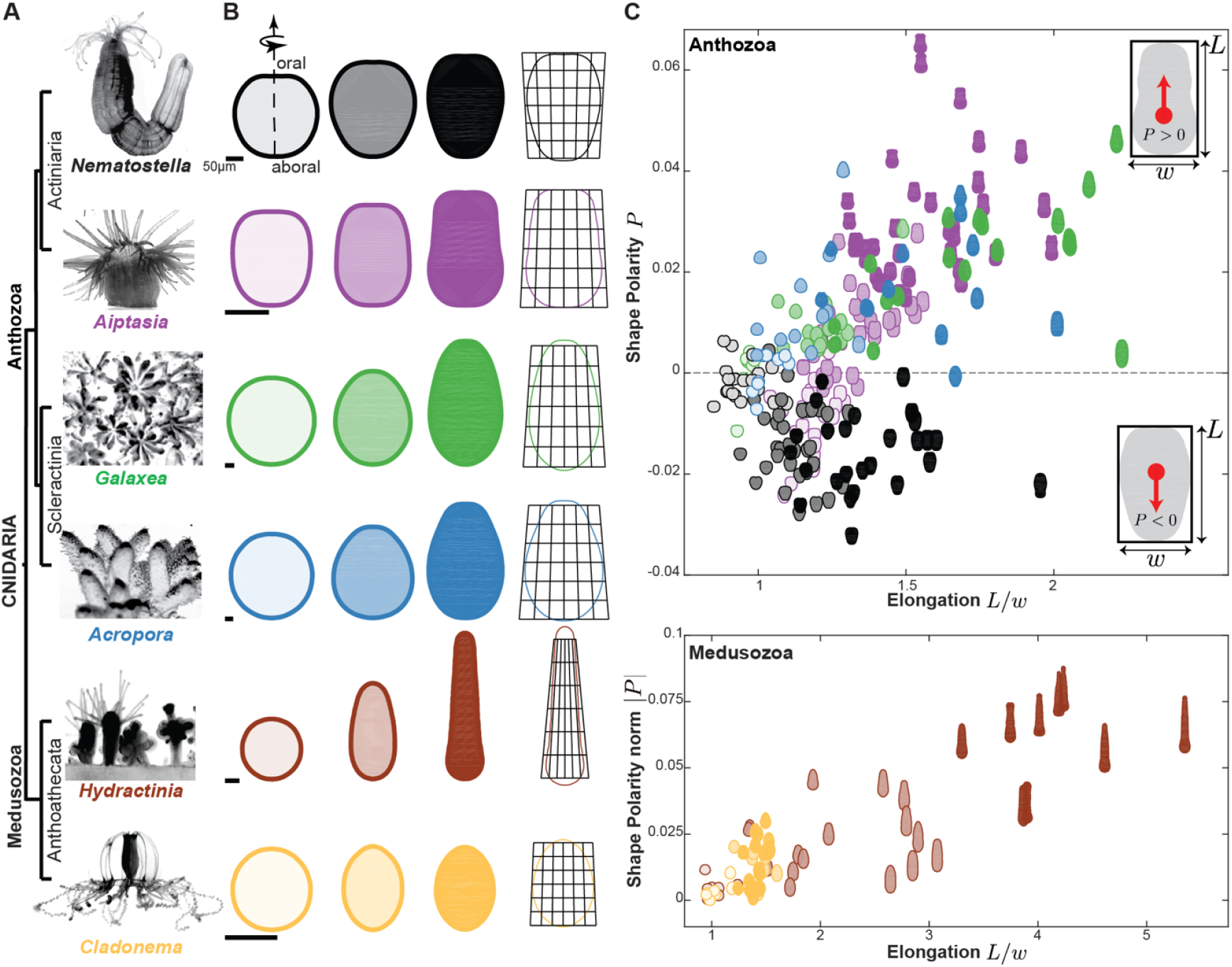
Diversity of cnidarian shapes from embryonic to planula stages. **(A)** A simplified phylogenetic tree of Cnidaria, incorporating subphyla and orders of the species examined in this study. Images shown are that of young adults in their sexual form. **(B)** (Left) Averaged shapes of three developmental stages: embryo, early planula, and planula (see Table S1) obtained from segmented bright field images. All six species transition from relatively spherical embryos to elongated planula stages. (Right) Planula shape outlines overlayed with “Thompsonian” affine transformation grids (Material and Methods). Scale bars: 50µm. **(C)** Morphospace of cnidarian larval shapes, positioned by their degree of elongation and shape polarity for Anthozoans (Top) and by absolute shape polarity for Medusozoans (Bottom). Segmented shapes are obtained from bright-field images. The red arrow indicates the shape polarity (P) along the oral-aboral axis.

While significant size differences were noted among species (fig. S1B), our focus was on shape variation. For each species, we mapped the outline coordinates of the organism from bright-field images across various developmental stages (Fig 1B, fig. S1A and Table S1). From these morphological outlines, we derived two main parameters to define a morphospace for shape diversity: aspect ratio (*L*/*w*) and shape polarity (*P*) (Fig. 1C, fig. S2A-B). The aspect ratio quantifies the degree of axial elongation by measuring the length-to-width ratio of the organism, while shape polarity measures the degree of shape asymmetry along the oral-aboral axis (Material and Methods). All embryos began with a relatively spherical shape and underwent elongation along their oral-aboral axis, progressively acquiring a polarized shape (Fig. 1C). The extent of elongation varied across species, with *Hydractinia* showing the most pronounced elongation. For shape polarity, *Nematostella* embryos broaden towards the oral pole during development (*P*<*0*). In contrast, the other anthozoans and *Hydractinia* developed a wider aboral pole (*P*>*0*), with varying degrees of positive shape polarity. These morphological parameters—aspect ratio and shape polarity—provide a comprehensive framework to quantify the variability in planula shapes. The holistic nature of morphological diversity was also illustrated by “Thompsonian” affine transformations, which effectively highlights shape differences between species (Fig 1B, Material and Methods) (*22*).

### Patterns of tissue architecture and myosin distribution

To explore the relationship between the observed diversity in larval shape and the mechanical properties of the ectoderm and endoderm across species, we analyzed the distribution of phosphomyosin light chain (pMLC) (fig. S3) as a proxy for contractile forces generated within the epithelium (*23*–*26*). In larvae of each species, pMLC intensity was enriched at the apical surface of the ectoderm and at the basal interface between the ectoderm and the endoderm (Fig 2A, fig. S2E). Notably, pMLC intensity was higher towards the aboral pole on the apical ectodermal surface, while the basal surface showed a more uniform and lower signal, except near the oral region in *Nematostella, Aiptasia*, and *Hydractinia* (Fig. 2B). We also noticed an ectodermal thickness gradient, with greater thickness at the aboral pole decreasing towards the lateral domain (Fig. 2A, fig. S2C). Additionally, the pMLC signal displayed an anisotropic distribution arising from planar alignment of stress fibers on the basal surface and cell outlines on the apical surface (Fig. 2C). To quantify alignment of stress fibers on the basal surface and cell outlines on the apical surface, we obtained from pMLC images the nematic order parameter, using a Fourier-transform-based pipeline (*27, 28*) (Fig. 2C, fig. S2D, fig. S3C; see Method). Specifically, *q* > *0* indicates circumferential order, *q* < *0* indicates longitudinal order and *q*=*0* indicates a lack of order. On the apical surface, the nematic order was variable: medusozoan planulae showed high circumferential order (*q* close to 0.5), while anthozoans exhibited a wide range of order magnitudes. In contrast, the basal surface consistently showed high circumferential order (*q* close to 0.5) across species, except in *Cladonema* (Fig. 1C). The profile of basal pMLC intensity further defined a boundary at the oral opening where the signal sharply decreased (Fig. 2A, fig. S4A) with position and opening angle that varied markedly among anthozoan species. Consistent with these differences, *Nematostella* exhibited extreme oral tissue involution (fig. S4), coral larvae showed moderate involution (*29*), and *Aiptasia* had none (fig. S4).

**Fig. 2.**
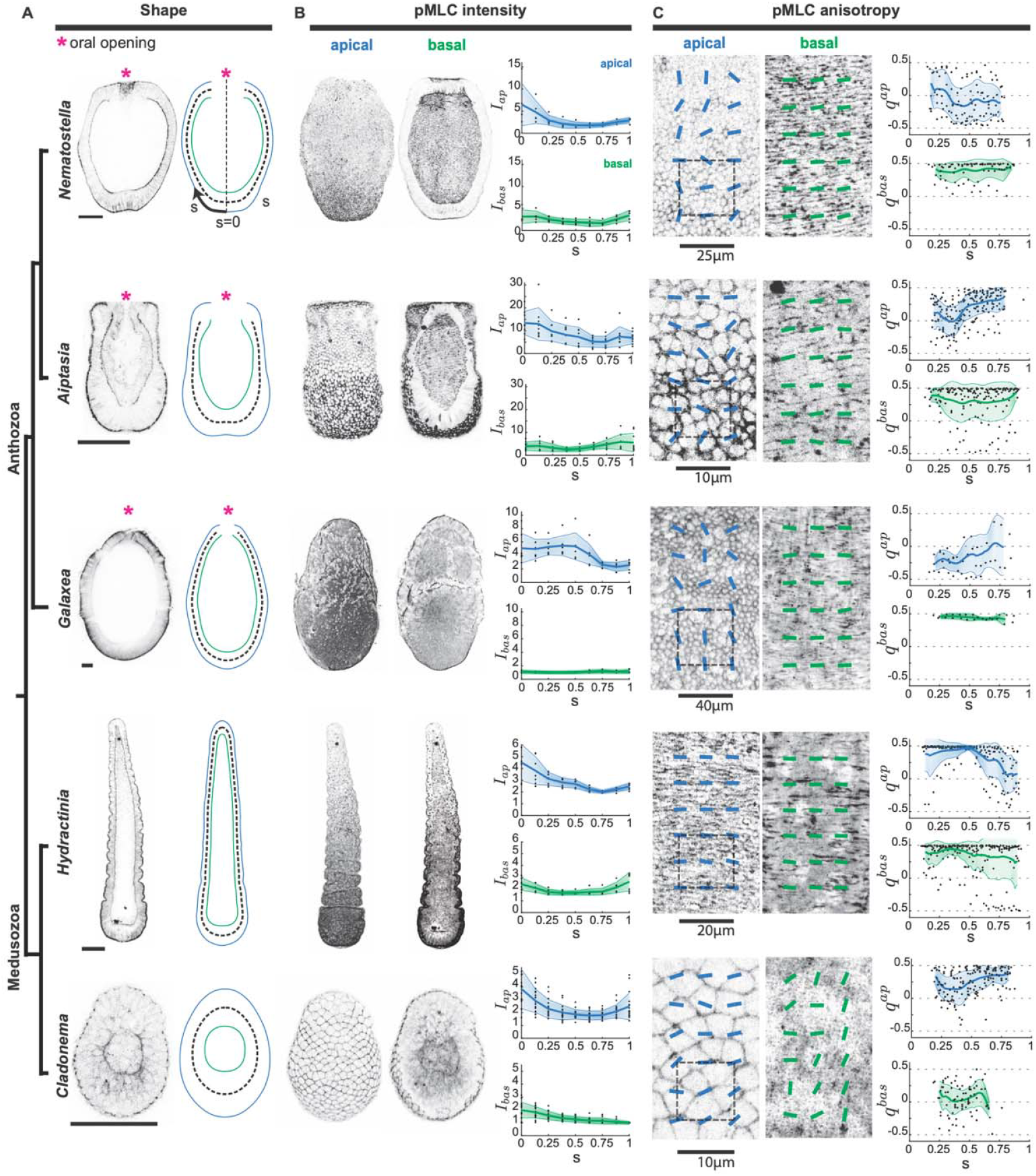
pMLC geometry, intensity and anisotropy in planulae. **(A)** (Left) Cross-section images of planulae stained for pMLC to visualize contractile structures. (Right) Representation of outer (apical ectoderm) surface, mean surface (dashed line) and inner (basal ectoderm/endoderm interface) surface contours averaged across the planula stage. denotes the normalized arclength coordinate along the oral-aboral axis. Asterisks mark the presence of an oral opening in anthozoans. Scale bars: 50µm. **(B)** Analysis of pMLC intensity patterns. (Left) Images of pMLC immunostainings showing apical and basal surfaces (same scale as in A). (Right) Profiles of pMLC intensity within the apical (top) and basal (bottom) surfaces as a function of normalized arclength coordinate (see Material and Methods). **(C)** pMLC anisotropy. (Left) Zoom-in views of apical and basal surfaces with superimposed bars representing anisotropy orientation, derived from a Fourier-transform analysis. Indicated scale bars correspond to the size of interrogation boxes used for each species (see Material and Methods). (Right) Profiles of nematic order parameter in the apical (top) and basal (bottom) surfaces as a function of normalized arclength coordinate. Shaded areas indicate standard deviations.

Despite the morphological diversity among cnidarian larvae, our comparative analysis suggests that underlying patterns of myosin and tissue architecture are surprisingly conserved at the mesoscale level, albeit with species-specific variations. These patterns are reflected in several supracellular features: a uniform circumferential nematic order on the basal surface of the bilayer, an excess of pMLC on the apical surface of the ectoderm accompanied by increased ectodermal thickness at the aboral pole, and a variable oral boundary geometry.

### Active surface model of shape diversity

To translate these supracellular patterns into biomechanical modules underlying larval shape diversity, we used an active surface model (*30*) adapted to the tissue architecture and axial symmetry of cnidarians (Fig. 3A, SI). This model represents the developing planula as an active viscoelastic shell. Forces within the surface originate from several contributions. Passive tensions and bending moments arise from surface area elasticity and bending rigidity, while active tensions and bending moments stem from myosin-dependent active tensions in the apical surface of the ectoderm and at the basal ectoderm/endoderm interface. We assume that contractile tensions are isotropic on the apical surface, acting uniformly in all directions. On the basal surface, we assume that actomyosin fibers generate a combination of isotropic and anisotropic surface tensions, according to their orientation (SI). For a given pattern of active tension and bending moments, the model predicts the shape of the mean surface, which averages the positions of the apical and basal surfaces through the middle of the ectoderm (Fig. 2A, Fig. 3A).

**Fig. 3.**
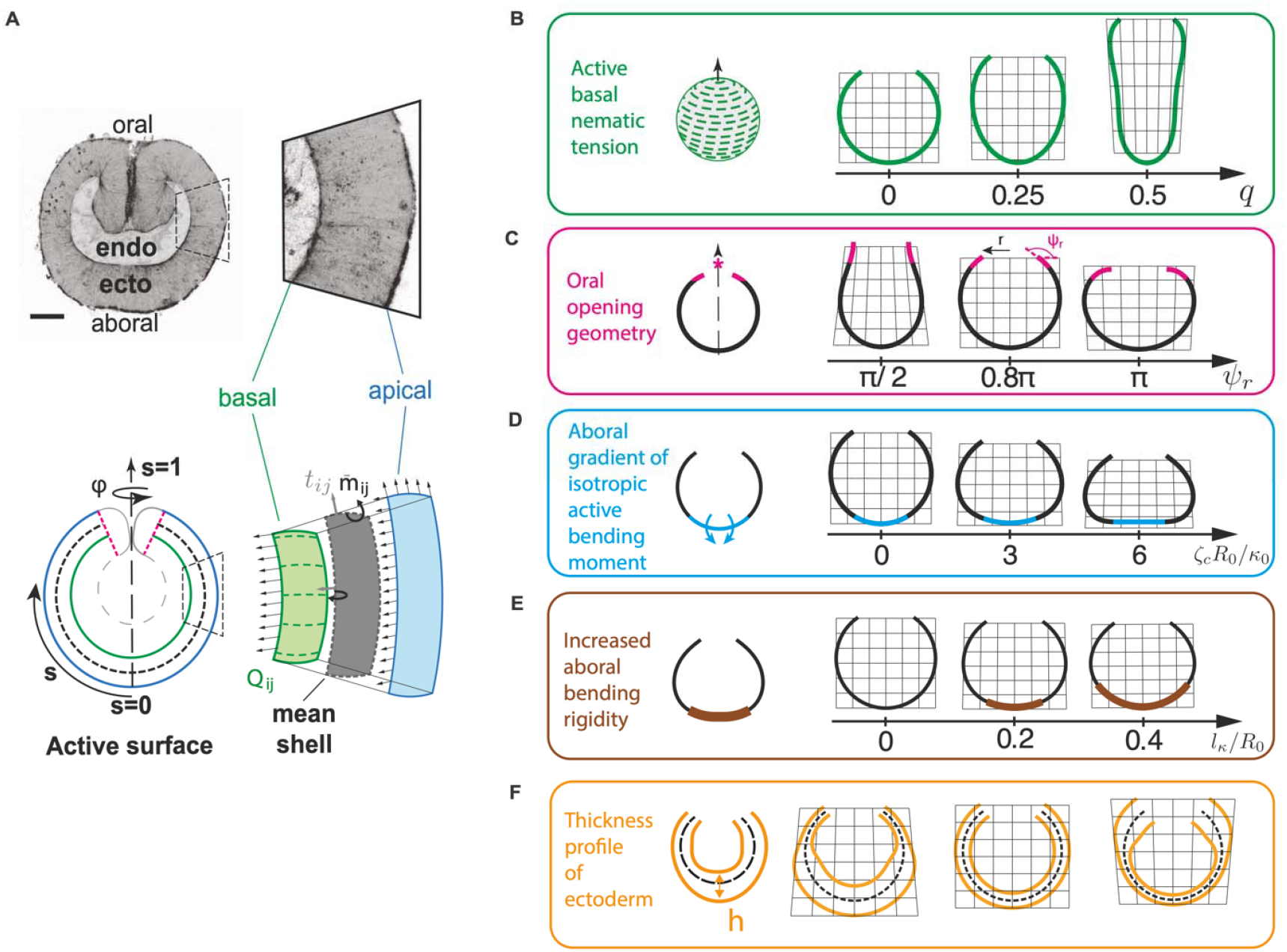
Inferred mechanical modules of shape variation. **(A)** (Top) Cross-section view of a *Nematostella* gastrula, illustrating the diploblastic organization with distinct ectoderm (ecto) and endoderm (endo) layers, and showing tissue apical and basal surfaces. Scale bar: 50µm. (Bottom) Schematics of active surface model., are the coordinates on the axisymmetric surface, the tensor of surface tension, the bending moment tensor, the tensor of nematic order. **(B-F)** Biomechanical modules underlying tissue elongation and shape polarity. Shapes correspond to steady-state simulations of the active surface model. Modules correspond to variations in basal nematic order and the resulting basal active nematic tension **(B)**, oral geometry (here angle at oral boundary) **(C)**, aboral active isotropic bending moment **(D)**, increased aboral bending rigidity **(E)** and ectoderm thickness **(F)**. is a reference radius, a reference bending modulus. See SI section 3.2 for other model parameters.

Using this theoretical framework, we investigated which basic physical modules, in the form of supracellular patterns of active tension, material properties and boundary conditions, could drive planula morphogenesis and shape diversification. We simulated the effects of such physical modules, starting from an open or closed spherical shape that mimics the gastrula (Fig. 3B-F, Fig. S4C). Active contractile tension directed by the nematic order *q* at the basal interface leads to shape elongation along the oral-aboral axis, which increases with the degree of order (Fig. 3B, movie S1). To explore the differences in the oral geometry observed in anthozoan planulae, we varied the angle of the oral extremities *ψ*_*r*_ of the mean surface, which influenced both shape polarity and aspect ratio (Fig. 3C, movie S2). As measurements of pMLC profiles showed an increase apical intensity on the aboral side (Fig. 2B), we tested for the effect of this pattern in the model. By increasing apical isotropic surface stresses on the aboral side, corresponding to an isotropic active bending moment of magnitude *ζ*_*c*_, we observed aboral shape flattening (Fig. 3D, movie S3). We also considered the possibility that increased thickness on the aboral side led to higher tissue bending rigidity. This resulted in aboral flattening for various domain size *l*_*k*_ of increased rigidity (Fig. 3E, movie S4). In addition, species-specific thickness profiles *h* modulate the shape polarity of the outer, apical ectoderm surface of the larva while keeping the mean surface constant (Fig. 3F, movie S5).

Collectively, these simulations predict that planula shape diversity can be driven by a combination of five mechanical modules: basal active nematic tension associated to basal nematic order, variations in oral geometry, active bending moments originating from varying apical contractility, regional differences in tissue bending rigidity and the outer, apical surface shape associated to gradients of tissue thickness.

### Developmental dynamics of planula shape

We then asked if the active surface model could account for the sequence of developmental shape changes by focusing on *Nematostella* (Fig. 1C). We performed pMLC staining at gastrula, early planula, and planula (Fig. 4A, fig. S5), and showed that an increase in the nematic order *q* on the basal ectoderm/endoderm interface was concurrent with an increase in aspect ratio (Fig 4A-B, fig. S5C). To model the morphogenesis of *Nematostella* planula, we imposed the boundary angle *ψ*_*r*_ and boundary radius *r* at the oral boundary of the tissue. We assumed that surface tensions at the apical and basal layers increase linearly with the normalized apical and basal pMLC intensities *I*_*ap*_ and *I*_*bas*_ with proportionality coefficients *β* (apical isotropic surface stresses) and *β*_*n*_ (basal anisotropic surface stresses). In other words, *β* and *β*_*n*_, which could be species-specific, allow to convert measured fluorescence intensities of pMLC into active contractile surface tensions. *β* effectively affects the magnitude of active bending moments and *β*_*n*_ the magnitude of nematic active tensions and bending moments (SI). We also allowed the bending rigidity to increase on the aboral side (by a factor *k*_*ab*_/*k*_0_ when the thickness was above 90% of its average. We found that with this model, a single non-zero value of *β*_*n*_, together with *β* = 0 and *k*_*ab*_/*k*_0_ =1, was sufficient to account for the gastrula, early planula and late planula shapes of *Nematostella* (Fig. 4C, SI). Overall, in the model, the progressive axial elongation was then explained by an increase in active nematic tension originating in the basal endoderm (Fig. 4C, fig. S5C).

**Fig. 4.**
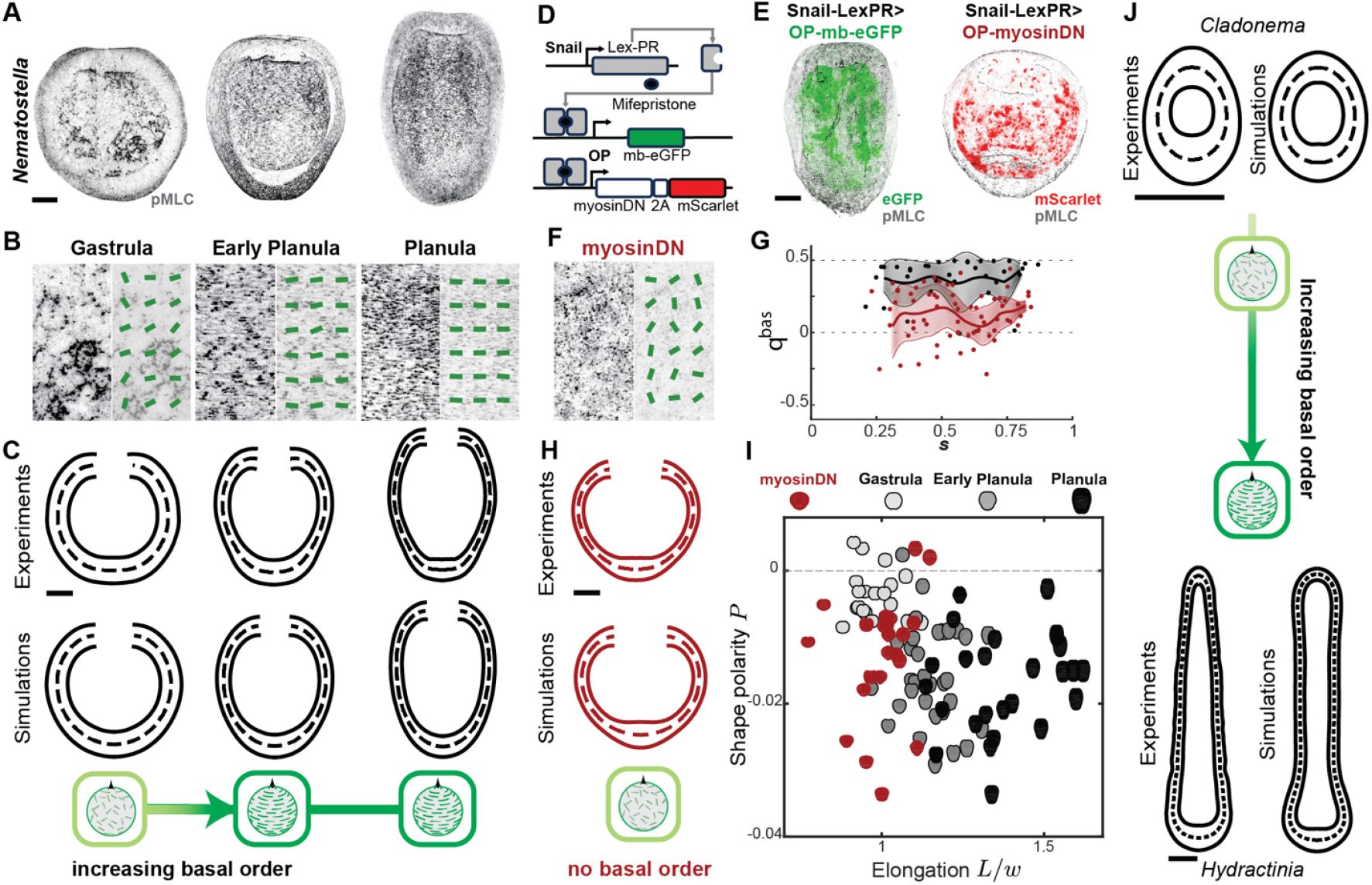
Basal nematic order drives axial elongation. **(A)** Images of pMLC immunostainings depicting the basal structures of developing *Nematostella*. **(B)** Zoom-in views with superimposed anisotropy bars, showing an increase of order after the gastrula stage. **(C)** (Top) Average experimental shapes of developing *Nematostella* and (Bottom) simulated shapes derived from implementing pMLC data into the active surface model. See SI section 3 for model parameters. **(D)** Cartoon of inducible genetic system. (**E**) (Left) Representative confocal image of an induced control animal carrying *LexPR*>*OP-mb-eGFP*, showing membrane-localized eGFP (mb-eGFP) expression in the endoderm, combined with pMLC staining. (Right) Representative confocal image of an induced animal carrying *LexPR*>*OP-myosinDN-2A-mScarlet*, showing activation of myosinDN in the endoderm, as indicated by mScarlet expression, combined with pMLC staining. (**F)** Detailed view of basal pMLC signal with anisotropy bars in myosinDN-induced *Nematostella*, indicating a loss of order in induced animals compared to wild-type. **(G)** Profiles of basal nematic order parameter as a function of normalized arclength coordinate, in control (black) and myosinDN-induced (red) animals. Shaded area indicate standard deviations. **(H)** (Top) Average experimental shapes of myosinDN-induced *Nematostella* and (bottom) simulated shape with active surface model. **(I)** Shape diagram mapping the elongation and shape polarity trajectories of developing *Nematostella* over three stages and highlighting the loss of elongation of induced animals expressing myosinDN. Shapes are obtained from bright field and pMLC staining images. **(J)** (Left) Average experimental shapes and (Right) active surface simulations for *Cladonema* and *Hydractinia*, with *Hydractinia* showing increased elongation and nematic order compared to *Cladonema*. See SI section 3 for model parameters. Scale bars: 50µm.

### Basal nematic order drives planula elongation

The model predicted that blocking basal contractility should interfere with axial elongation. Testing this hypothesis requires a method to selectively disrupt tissue contractility in a tissue-specific manner. We therefore developed a chemically inducible gene expression system in *Nematostella* (Fig. 4D, fig. S6A) (*31*), which we used to drive a dominant negative version of myosin (myosinDN) (*32*) specifically in the endoderm. This system uses the Lex-PR transactivator, which is responsive to Mifepristone (fig. S6A). Spatial specificity was achieved using the *snail* promoter (*33, 34*), which drives expression exclusively in the endoderm. Temporal control was provided by the addition of Mifepristone, inducing the expression of myosinDN (Fig. 4D).

We established a LexPR transgenic line with the Snail promoter (fig S6A-B). This Snail-LexPR driver line was crossed with an effector line carrying an inducible transgene for myosinDN. Upon induction of myosinDN, we observed a marked reduction in basal pMLC intensity and nematic order (Fig. 4E-G, fig. S6C). Surprisingly, in control planulae, the basal myosin signal and oriented fibers appeared to come mostly from the ectoderm (fig. S6A). As the perturbation is specific to the endoderm, this indicates that basal nematic order originates from a cross-talk between the endodermal and ectodermal layers. Furthermore, this perturbation did not significantly affect oral geometry, ectodermal thickness, and apical myosin profiles, suggesting a precise disruption of the basal interface organisation (fig. S6C-F). Crucially, as predicted, animals expressing myosinDN did not undergo axial elongation (Fig. 4H, I). Instead, these DN-myosin-induced animals showed a round, slightly polarized shape (Fig. 4H, I). The model also predicted well this modified shape using the perturbed profiles of pMLC intensity and order (Fig. 4H, fig. S6C-F). These results demonstrate that ordered supracellular myosin activity at the basal interface is essential for axial elongation.

Building on this finding, we asked if this relationship might be evolutionarily conserved. All anthozoan species showed high basal order together with axial elongation (Fig. 2A, C). However, our quantification of nematic order in medusozoans indicated that *Cladonema* has almost no nematic basal order, in sharp contrast to *Hydractinia* (Fig. 2C). Taking into account this difference, together with the closed oral shape of these species, the model could capture the weak elongation of *Cladonema* and strong elongation of *Hydractinia* (Fig. 4J, fig. S7A). Altogether, these results indicate that species with high basal nematic order utilize this mechanical module for larval elongation, supporting the hypothesis that basal nematic order resulting in basal active nematic tension is a key driver of axial elongation conserved across cnidarians.

### Oral geometry constraints anthozoan shape polarity

To uncover the biophysical factors that govern shape polarity, we performed an in-depth comparative analysis of the oral geometry in anthozoans (Fig. 5A), as this module had the most obvious effect on shape polarity in simulations (Fig. 3C). The boundary angle *ψ*_*r*_ and relative radius *r*/*R*_0_ at the oral boundary varied significantly between species (Fig. 5B). *Nematostella*, for instance, showed a small, nearly horizontal boundary (*ψ*_*r*_/ *π*) and *Aiptasia* a nearly vertical, large boundary 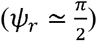. The angle at the oral boundary remained relatively constant throughout the transition from gastrula to planula in *Nematostella* and *Aiptasia* (fig. S8A). We then asked to what extent shape polarity could be predicted solely from the oral geometry in the model. We calculated the predicted shapes for varying values of boundary angle and radius of the oral boundary, with no active tension or bending moment (Fig. 5B). We noticed that simulated shapes exhibit negative polarities along the descending diagonal in the space (*ψ*_*r*,_ *r*/*R*_0_), and positive polarities away from the diagonal (Fig. 5B, fig. S8B). We observed a similar pattern in experimental data, suggesting a role for oral geometry in setting shape polarity (Fig. 5B). Despite this broad agreement, the polarity of *Aiptasia* was more negative than expected purely based on oral geometry, indicating a role of other modules.

**Fig. 5.**
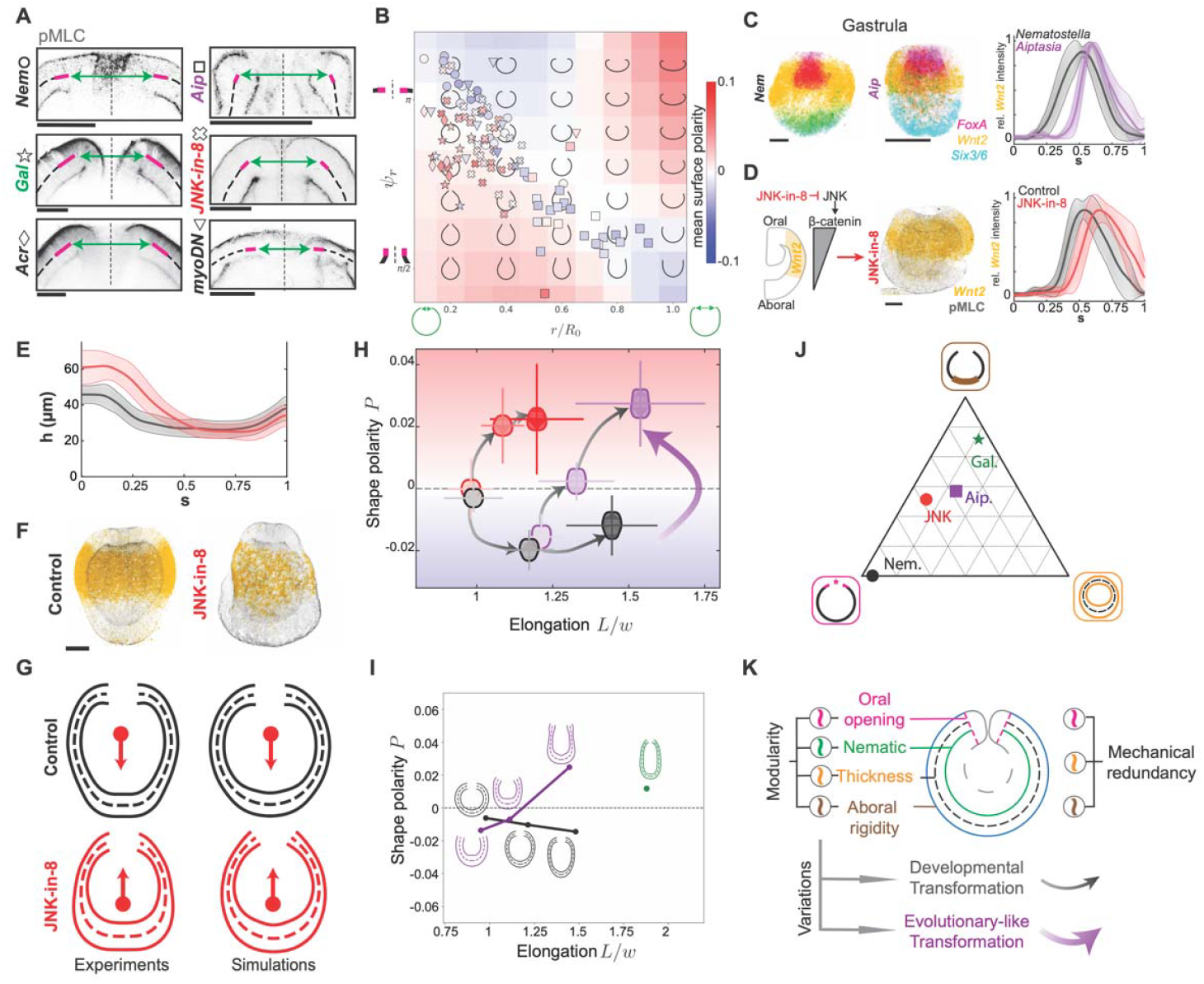
Biomechanical determinants of shape polarity. **(A)** Zoom-in views of pMLC immunostainings showing the oral opening of anthozoan wild-type planulae and *Nematostella* perturbations. Dashed lines represent the mean surfaces, solid lines and double arrows indicate angles and diameters at the oral boundary. **(B)** Oral module shape diagrams. Black shapes and background color correspond to passive surface simulation results as a function of the normalized oral radius and angle. Symbols correspond to individual larvae at all stages, positioned in the diagram according to experimentally measured values of and, colored according to measured shape polarity, with symbol shapes corresponding to species and treatments as indicated in A. **(C)** Images of HCR stainings highlighting conserved FoxA, Wnt2 and Six3/6 domains that pattern the oral-aboral axis in *Nematostella* (Left) and *Aiptasia* (Middle). Profiles of relative Wnt2 expression along the aboral-oral axis at the gastrula stage, in *Nematostella* VS *Aiptasia* (Right). **(D) (**Left) Cartoon illustrating the effect of the JNK-in-8 treatment on *Nematostella*’s axial patterning. (Middle) Sum projections of pMLC immunostainings combined with Wnt2 HCR in JNK-in-8-treated *Nematostella* at the gastrula stage. (Right) Profiles of relative Wnt2 expression along the aboral-oral axis at the gastrula stage in control and JNK-in-8 treated *Nematostella*. **(E)** Profiles of ectodermal thickness as a function of normalized arclength coordinate at planula stage (control, black; JNK-in-8, red). (**F**) Sum projections of pMLC immunostainings combined with Wnt2 HCR in control and JNK-in-8-treated *Nematostella* at the planula stages. **(G)** Average experimental shapes of JNK-in-8-treated *Nematostella* and corresponding simulated shape, at planula stage (red). Black shapes correspond to control average experimental shape and simulated shapes at planula stage. (**H**) Morphospace of larval shapes of developing *Nematostella* (wild type, black; JNK-in-8, red) and *Aiptasia* (purple), positioned according to outer surface elongation and shape polarity. Black arrows indicate developmental trajectories, purple arrow evolutionary-like transformations. Shapes are obtained from pMLC staining images. (**I**) Morphospace of simulated larval shapes of *Nematostella, Aiptasia* and *Galaxea*, positioned according to outer surface elongation and shape polarity. (**J**) “Mechanical redundancy triangle”, showing the relative weight of three modules (increased aboral bending rigidity, oral geometry, thickness profile) in determining the simulated shape polarity of indicated species, at late planula stage. (**K**) Variations in biomechanical modules account for both developmental and evolutionary-like transformations. Scale bars: 50µm. Shaded areas indicate standard deviations. G,I, J: See SI section 3 for model parameters.

### Perturbing axially organized modules reverses polarity

To further explore the relationship between mechanical modules and shape polarity, we thought to perturb axial patterning, since these modules are organized along the oral-aboral axis, which is the primary body polarity of cnidarians. In *Nematostella*, early embryonic patterning relies on beta-catenin signaling (*35*), delineating three ectodermal expression domains: Six3/6 at the aboral pole, Wnt2 in the mid-domain, and FoxA in the involuting oral tissues (Fig. 5C, fig. S9A). Using hybridization chain reaction (HCR (*36*)), we also identified these morphogenetic territories are conserved in *Aiptasia* (Fig. 5C), but their spatial distribution varies between the two species (fig. S9A). Notably, the *Nematostella* Wnt2 expression pattern expanded closer to the aboral pole, whereas in *Aiptasia*, it extended more toward the oral pole (Fig. 5C, fig. S9A). We hypothesized that this divergence underlies the evolutionary differences in spatially patterned modules and their effects on shape polarity.

Following these observations, we experimentally tested whether modifying the early axial patterning in *Nematostella* to mirror that seen in *Aiptasia* would affect shape polarity (Fig. 5D). The inhibition of JNK signaling, reported to modulate beta-catenin activity (*37*), induced a clear shift in the Wnt2 expression domain in embryos, closely resembling the pattern observed in *Aiptasia* (Fig. 5D). Measuring the oral geometry in JNK-inhibited *Nematostella* larvae showed reduced boundary angle *ψ*_*r*_, bringing *Nematostella* oral geometry closer to *Galaxea* rather than *Aiptasia* (Fig. 5A, B). At the same time, the thick aboral ectoderm domain became thicker and broader, making it more similar to *Aiptasia* (Fig. 5E, fig. S2C). Additionally, the aboral domain of higher apical pMLC increased in size and intensity (fig. S9B-C), while basal nematic order remained circumferential (fig. S9B-C). Strikingly, JNK-inhibited *Nematostella* larvae were elongating normally but with inverted polarity (Fig. 5F, G), shifting *Nematostella* position in the morphospace closer to *Aiptasia* (Fig. 5H). Taking measured changes in oral geometry, nematic order, tissue thickness and apical pMLC intensity into account in the active surface model, we could explain the reversal of shape polarity following JNK treatment, provided that the increased bending rigidity module (*k*_*ab*_/*k*_0_ >1 but not the active isotropic bending moment module (*β* = 0), acts on the aboral side of wild-type and JNK-treated *Nematostella* (Fig. 5F, fig. S9D-F, SI). Thus, manipulating mechanical modules recapitulates evolutionary-like shape transformation by modifying developmental trajectories (Fig. 5H, fig. S9F).

### Mechanical redundancy in shape determination

We then tested if active surface model simulations could also recapitulate shapes of *Aiptasia* and *Galaxea* by adjusting the coefficients *k*_*ab*_/*k*_0_ and *β*_*n*_ setting *β* = 0, and using experimental measurements of pMLC intensity, nematic order, oral geometry and ectoderm thickness (Fig.2, fig. S5). We found that this was the case for *Aiptasia* and for *Galaxea*, with close values of the adjusted coefficients *k*_*ab*_/*k*_0_ and *βn* between *Aiptasia* and *Nematostella*, two species of sea anemones (Fig. 5I, fig. S7, SI). Notably, the model could account for the experimentally observed switch of shape polarity of *Aiptasia* between early and late planula (Fig. 5H, 5I, fig. S7). To then determine the relative contributions of the three modules of increased aboral bending rigidity, oral geometry and ectoderm thickness to shape polarity, we performed simulations where these modules were “turned off” (Fig. 5J, fig. S10). The model indicated that the oral geometry module was largely responsible for *Nematostella* shape polarity, the increased aboral bending rigidity module for *Galaxea* shape polarity, while both modules contributed to define the shape polarity of *Aiptasia* and JNK-treated *Nematostella* (Fig. 5J). Thus, shape polarity is not determined by a single module across species, but by a combination of modules that differ between species, highlighting the redundancy of mechanical modules for shape determination (Fig. 5J).

## DISCUSSION

Our study establishes a framework for evo-devo-mechanobiology (*38*) by demonstrating the critical role of biomechanical modules in shaping cnidarian larval development and diversity. We define cross-species biomechanical modules as conserved supracellular units that direct and constrain morphogenesis. These modules serve as dynamic building blocks of morphogenesis, capable of changing across both developmental and evolutionary time scales (Fig. 5K). Here, we have identified such modules across a set of cnidarian larvae. The nematic order of basal stress fibers emerged as the main module driving axial elongation in both anthozoans and hydrozoans. In contrast, shape polarity arises from distinct combinations of two main modules—oral geometry and spatially patterned bending rigidity—with a minor contribution from tissue thickness in *Aiptasia*. In anthozoans, species-specific variations in oral geometry, likely set during gastrulation (*39*), establish boundary conditions for morphogenetic processes that ultimately influence organismal form, indicating a potential role of gastrulation modes in subsequent shape determination.

Experimental manipulations of these supracellular modules in *Nematostella* demonstrated their causal role in morphogenesis. Reversing the shape polarity of *Nematostella* showed that variations in these modules drive shape diversity. Furthermore, the ability to produce similar shapes through different mechanical module combinations also suggests a form of mechanical redundancy (Fig. 5K), analogous to genetic redundancy (*40, 41*), where multiple genes can fulfill similar functions. Such mechanical redundancy might provide resilience against mutations and enable evolutionary innovation, allowing diverse morphologies to evolve without disrupting fundamental developmental processes. We demonstrated this concept by manipulating axial patterning-driven mechanical modules in *Nematostella*, successfully shifting its larval shape closer to *Aiptasia* within the morphospace, with a combination of features present in both *Galaxea* (oral geometry) and *Aiptasia* (aboral thickness).

While we focused on specific mechanical modules, other factors likely contribute to larval shape. The polarized shape of *Hydractinia*, for instance, hints at additional determinants not captured in our model (Fig. 4J). Further exploration into the developmental origins of nematic order, potentially linked to the planar cell polarity pathway (*42*), will deepen our understanding of mechanical module establishment. Species-specific oral geometry may also reflect adaptations to distinct feeding strategies and environmental interactions, such as the predatory behavior observed in *Aiptasia* larvae (*43*). In summary, we propose that shape diversity can be explained at three distinct levels of biological organization: (*i*) the microscopic level, involving numerous molecular and cellular parameters; (*ii*) the mesoscopic level (*44*), where many of these parameters converge into a few physical parameters defining mesoscale biomechanical modules; and (*iii*) the macroscopic level, where variations in these modules generate observable shape diversity. This framework opens new avenues to explore the evolutionary pressures on these mechanical modules and the processes that drive morphological diversity and innovation.

## Acknowledgments

We thank Kresimir Crnokic for his support in animal husbandry. We are thankful to Rashmi Priya and Héctor Sánchez-Iranzo for sharing the plasmids for the inducible system, and to Marie Anzo for cloning the Snail promoter into the driver plasmid. We thank the Ikmi and Salbreux labs for their comments on the work. Additionally, we also thank Steffen Lemke, Marie Manceau and Jean-Paul Vincent for their comments on the manuscript.

## Author Contributions

I conceived the initial idea for the project. RB, NC, GS, and AI conceptualized the work. RB and AI designed the initial experiments with the support of ADM and GS. The initial theoretical model was designed by DK and GS. NC and GS integrated the biophysical data into the theoretical model. NC refined the model and ran all the simulations. RB conducted most of the experiments by performing the pharmacological treatments, induction experiments, stainings, imaging, and data quantification. SB performed two-photon imaging of corals. PJS helped with staining and confocal imaging. RB established and performed HCR. AI designed and injected the constructs of the inducible system. GB and PC cloned and tested the inducible system. AI and RB selected and grew the driver and effector lines. RB performed the genetic crosses. SR and AG provided live *Aiptasia*. JC provided fixed *Acropora*. NS and PC provided fixed *Galaxea*. CC provided live *Hydractina*. SF and YN provided fixed *Cladonema*. RB, NC, GS and AI analyzed the data. RB and NC generated the figures. GS and AI drafted the manuscript with the support of RB and NC. All authors reviewed the manuscript.

## Data and materials availability

All data needed to evaluate the conclusions are present in the paper and the supplementary materials. Simulation codes are deposited at https://github.com/DianaKhoromskaia/EpithelialShell. Transgenic lines and plasmids are available upon request.

## Materials and methods

### Animals

*Nematostella* adults were maintained with 12 parts per thousand (ppt) artificial seawater (ASW; sea salt, instant ocean) at 17°C in the dark. Spawning was induced by putting the animals overnight in a box with white light (6-8hours), raising the temperature to around 28°C (*44*). Spawning occurred within 3–4 h after a cold (17°C) water change. Eggs were de-jellied in 4% cysteine solution (Sigma, 168149) in ASW for 9 minutes, then washed with ASW three times and fertilized. *Aiptasia* adults were maintained with 36ppt ASW at 26°C with a “night/day” cycle, using a white light from 8pm to 8am. A 29-days lunar cycle was simulated by increasing the incubator temperature to 29°C and by illuminating during the “night” with blue light for 5 days - day 2 to day 6 of the cycle (*45*). Spawning occurred at ∼1-2pm for several days around two weeks after induction with the blue light (eggs and sperm have been collected at various amounts from day 15 to day 25). Animals were put back in resting mode at 26°C without blue light after two spawning cycles. *Hydractinia* colonies were cultured with 29ppt ASW at 19°C in the dark. Spawning occurred about 90mins after taking the colonies out of the dark (*20*). Other species were maintained, fixed and dehydrated in host laboratories: respectively, Carnegie Science (USA) for *Galaxea*, Horniman Museum and Gardens (UK) for *Acropora*, and Graduate School of Pharmaceutical Sciences (University of Tokyo, Japan) for *Cladonema*), and then shipped to EMBL Heidelberg where stainings were performed. Fixation and dehydration methods are presented in Table S2.

### Stainings

Animals were rehydrated by serial washes in ethanol or methanol solutions diluted in PTw (PBS with 0,1% of Tween20, Sigma, P1379), followed by five washes in PTw.

#### Immunohistochemistry (IHC)

Samples were blocked in a buffer containing PTx (PBS with 0.2% Triton, Sigma, T8787), 0.1% DMSO (Thermo Fisher, 85190), 1% BSA (Sigma, A2153), and 5% Goat serum (Sigma, G9023) for 1 hour at room temperature (RT). Animals were then incubated with primary antibodies in the blocking solution overnight (12-16 hours) at 4°C. Primary antibodies were anti-pMLC (*46*) (Rabbit; Cell signaling technology, 3671S, dilution 1:50), anti-GFP (Mouse; Thermo Fisher, A-11072, 1:500) and anti-RFP (Rat; Chromotek 5F8, 1:500). After washing 5 times with PTw, samples were incubated with their corresponding secondary antibodies. Secondary antibodies were goat-anti-rabbit Alexa-594 (Thermo Fisher, A-11072, 1:500), donkey-anti-mouse Alexa-488 (Thermo Fisher, A-21202, 1:500) and goat-anti-rat Alexa-647 (Abcam, Ab150159, 1:500) in PTx for 5-6 hours at 4°C, and washed again 5 times in PTw. pMLC immunostaining in *Acropora* captured oral geometry but not reliably myosin profiles.

#### Hybridization Chain Reaction (HCR)

HCR kits (version 3) containing DNA probe sets, amplifier sets, along with hybridization, amplification and wash buffers were purchased from Molecular Instruments (http://molecularinstruments.org) and protocols available in other species (*47*) were adjusted for

#### Nematostella and Aiptasia

Rehydrated samples were incubated in a 20µg/mL proteinase K (Promega, V3021) solution with PTw (10min for *Aiptasia*, 30min for *Nematostella*), followed by 2 washes in PTw and post-fixation in 4% PFA (paraformaldehyde, Electron Microscopy Sciences, E15710) with PTw. After five washes in PTw, animals were transferred in 5X SSCTw (SSC, Sigma, S6639 with 0,1% of Tween20) and pre-hybridized in probe hybridization buffer (ordered at Molecular Instruments) for 30min at 37°C. Samples were then incubated in probe solutions containing 3pmol of each gene-specific probe set overnight at 37°C. Probe sets were designed based on published sequences for each gene: For *Nematostella*, FoxA (AY457634.1), Wnt2 (AY725201.1) and Six3/6 (KC137590.1); For *Aiptasia*, FoxA (XM_028657059.1), Wnt2 (XM_021040219.2) and Six3/6 (XM_021061437.2). FoxA, Wnt2 and Six3/6 were designed with amplifiers B2, B3, and B1, respectively.

Animals were then washed 3 times in probe wash buffer (Molecular Instruments) for 20min at 37°C, and 2 times in 5X SSCTw for 5min at RT. In parallel, hairpin duos (B1-Alexa 488, B2-Alexa 546 and B3-Alexa 647) were separately heated for 90s at 95°C, cooled down at RT for 30min, and diluted 1:50 in probe amplification buffer. Washed animals were incubated in the hairpin solution overnight at RT in the dark. Finally, animals were washed 5 times in PTw.

HCR and IHC were combined by performing HCR first, followed by 3 washes in PTx 0,2%, and then IHC from the blocking step. Samples were protected from light in all steps following incubation in hairpin solutions (for HCR) or secondary antibody (for IHC).

### Pharmacological treatments

JNK-in-8 (Sigma, SML1246) solution was prepared by diluting 40mM stock solution (stored in 100% DMSO) dissolved with 12ppt ASW. For treatments, DMSO concentration was adjusted to 0,1% for both control and JNK-in-8 conditions. Animals were incubated with 20μM JNK-in-8 from 1hpf at 21°C and solutions were refreshed at around 42hpf. This drug was tested in three replicates. After fixation (see Table S1), controls and JNK-in-8 treated animals were stained with pMLC (IHC) and Wnt2 (HCR).

### The inducible system

The design of the inducible genetic system was developed based on the Zebrafish constructs (*31*) with optimization for *Nematostella* gene expression and transgenesis. A dual-plasmid framework comprising driver and effector plasmids was constructed. The backbones of these plasmids were based on the *Nematostella* SceI transgenesis plasmid (*48*).

The driver plasmid was engineered by cloning a multi-cloning site (MCS) upstream of the open reading frame for the artificial transcription factor (LexPR) representing the fusion of a DNA binding domain from the bacterial LexA protein, a truncated ligand binding domain from the human progesterone receptor and an activation domain from the human NF-κB/p65 protein, followed by an SV40 polyA signal. Additionally, a second cassette for a membrane-bound eGFP followed by an SV40 polyA signal was cloned in the same transgenesis plasmid and driven by the LexA operator. The Sce-I restriction sites flanked this composite construct to enable transgenesis (fig. S7A). To generate a driver plasmid for specific endodermal expression, the Snail promoter (*34*) was PCR-amplified from the genomic DNA and inserted into the driver plasmid linearized at the AflII site (Primers are listed in Table S3).

To generate the effector plasmid, a first construct containing a multi-cloning site (MCS) was cloned between the LexA operator and a 2A-mScarlet with SV40 polyA, followed by a second selection cassette encoding an eGFP with a nuclear localization signal (NLS) under the control of the actin promoter (*49*). Like the driver, this plasmid is bordered by SceI sites to facilitate transgenesis (fig. S7A). A dominant-negative allele of Myosin (myosinDN) was synthesized by mutating Threonine 18 to Alanine and Cysteine 19 to Alanine, as described in Priya et al (*32*). This myosinDN construct was amplified and cloned in the MCS of the effector plasmid in frame with the coding sequence of the 2A-mScarlet using AscI/AgeI restriction sites (Primers are listed in Table S3).

All PCR fragments were amplified with the Q5 HiFi DNA Polymerase KIT (#MO491) or the Thermo Scientific Phusion Green Hot Start II HiFi PCR Master Mix (#F-565S). Cloning was performed with the NEB Quick Ligation kit (#M2200) or the NEB Builder HiFi DNA Assembly Master Mix (#M5520). Sequencing confirmed all plasmid constructs.

### Transgenesis

Transgenic lines were established via meganuclease-mediated transgenesis (*48*). For the driver line, primary polyps displaying membrane bound-eGFP expression were identified through Mifepristone administration (1uM) in artificial seawater (12ppt) and selected as potential founders. These GFP-positive polyps were raised to sexual maturity and crossed with wild-type animals to derive F1 Snail-LexPR lines. F2 homozygotes were obtained by intercrossing F1 individuals from the same founder. Potential founders for the effector line were selected based on Act-NLS::eGFP expression, grown to sexual maturity matured, and crossed with wild-type counterparts to establish the F1. F2 homozygote animal for the effector line was obtained by intercrossing F1 individuals.

### Crosses and induction

F2 homozygotes Snail-LexPR driver lines were crossed with F2 homozygotes DN-myosin effector lines and fertilized eggs were incubated at 27°C. At 4hpf, the 12 ppt ASW was replaced with 1µM mifepristone diluted in 12 ppt ASW and this solution was refreshed at around 24hpf. On day three post-fertilization, both GFP and mScarlet signals were clearly visible, and animals showing broad endodermal induction were selected for analysis.

### Imaging

Stained animals were initially rinsed with PBS and subsequently cleared in 80% glycerol solutions. The samples were carefully transferred to microscope slides, and secured with thin layers of double-sided tape. Any excess glycerol was then removed before placing a glass coverslip over the samples for imaging. For the microscopy, a Leica SP8 confocal microscope was employed for all species except corals. Corals were imaged using a Zeiss LSM780 NLO two-photon microscope due to their large size. The specific objectives utilized for each species are detailed in Table S4. Additionally, for bright field imaging, a Leica MZ16F stereomicroscope equipped with a QImaging QICAM FAST 12-bit camera was used to capture the overall morphologies of all specimens.

### Image analysis

Microscopy images were oriented with the oral pole at the top and processed using both Fiji and MATLAB.

#### Surface coordinates and morphometrics

Animals captured in bright field images were automatically segmented using a custom MATLAB program (Fig. 1, S1, Fig. 4I, Fig. 5H). The process involved the following steps: Gaussian smoothing was applied using the MATLAB function ‘imgaussfilt’, followed by thresholding, and morphological operations like opening (‘imopen’) and closing (‘imclose’) with parameters tailored to each image’s properties to generate binary masks. The coordinates (*x,z*) from the outlines of these segmented masks were extracted using the ‘bwconncomp’ function. The data were organized into left (*x* < 0) and right (*x* >= 0) halves and interpolated into 101 evenly spaced points ranging from 0 (aboral pole) to 1 (oral pole) using the ‘interparc’ plugin (available at mathworks.com/matlabcentral/fileexchange/34874-interparc). This established a normalized arc-length coordinate *s* = 0,0.01, …, 1, which was utilized for averaging curves, and serving as a spatial reference for aligning subsequent parameters. For each contour, the coordinates of the left and right sides were averaged along the *s* coordinate to create a symmetrized shape (fig. S3A).

For pMLC IHC image segmentation (Fig. 2, Fig. 4. Fig. 5, Fig. S2, Fig. S5). apical and basal surfaces were manually outlined using the segmented line tool in Fiji from cross-sections. Symmetrized versions of these layers were generated similarly to the method described for bright field images. To calculate the average positions of the apical and basal surfaces, the mean surface was determined by computing the intersections of successive basal normal lines with the apical surface and extracting the midpoints of these line segments. The resulting (*x, z*) coordinates were interpolated into evenly spaced points as described previously. Here as well, average shapes between individuals of a given stage and species are obtained by averaging *x*(*s*),*z*(*s*) curves. Additionally, to determine the oral opening angle, a line was fitted using the ‘polyfit’ function (degree *n* = 1) over the last five points of the interpolated mean surface (fig. S2A). The ectodermal thickness *h* (s), measured as the distance between basal layer coordinates (*x*_*bas*_ (*s*), *z*_*bas*_ (*s*)) their closest point in the apical surface, was smoothed using a moving average with a window size equivalent to 5% of the curve length.

Average shapes between individuals of a given stage and species are obtained by averaging *x*(*s*),*z*(*s*) curves. The length *L*. defined as *L* = (*z*_*max*_ — *z*_*rnin*_) and width *w* = 2*x*_*max*_ were extracted to obtain the shape elongation. *L*/*w*. The shape polarity *P* was obtained by evaluating:

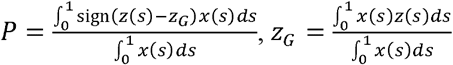

where *z*_*G*_ corresponds to the *z* position of the surface centroid, and P measures the relative surface area above and below the centroid; assuming axisymmetry of the shape.

Ill order to evaluate *ψ*_*r*_. and *r*/*R*_0_ (Fig. 5B. SI), we evaluate the boundary angle *ψ*_*r*_ and radius *r* at the oral pole, for individual or averaged shapes (Table S6). The reference radius *R*_0_ is set by evaluating the surface area *A* of cnidarian larvae of different species, and taking the radius of curvature of the spherical cap with same area:

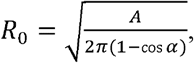

where the angle *α* =0.7*π* if the shape is open and *π* otherwise.

#### pMLC intensity

pMLC intensity profiles were extracted from orthogonal views using Fiji, by drawing lines from the apical surface into the basal surface around *s* = 0,0.125,0.250, …, 1, which were evaluated from segmentation of the contour of the shape (fig. S3B). At each position *s*, intensity profiles obtained were further processed using MATLAB, as follows. The curves were smoothed using a moving average of 1µm. Local maxima corresponding to the apical and basal pMLC-enriched surfaces were detected using the ‘findpeaks’ function in MATLAB. The intensity at these peaks, *M*_*ap*_ for the apical surface and *M*_*bas*_ for the basal layer, were normalized using the intensities *m*_*ap*_ and *m*_*bas*_ measured in the cytoplasm at a distance *D* from the peaks. Values of the distance *D* chosen for different species are listed in Table S5. We then defined the normalized intensities 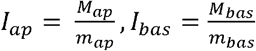 (fig. S3B). The value *I*_*bas*_ =1 was attributed when no basal peak was detected in specimens such as *Galaxea* and *Cladonema* planulae.

#### pMLC anisotropy

To quantify anisotropy, projections of the outer and inner layers were initially obtained using the Minimum Cost Z Surface Projection plugin available on FIJI (https://imagej.net/plugins/minimum-cost-z-surface-projection). Regions near the poles were cropped to minimize errors from animal curvature. A Fourier Transform (FT)-based method was then applied to these projections to assess coarse-grained anisotropy (*27*). Images were segmented into square interrogation boxes of uniform size, adjusted for each species, with a 50% overlap. The FT was computed on these boxes using MATLAB’s fast FT algorithm (fft2.m). Resulting Fourier space patterns were binarized by retaining only the 5% brightest pixels, yielding filled ellipses from which inertia matrices were computed and diagonalized. This allowed extracting a scalar alignment parameter *A*, defined as 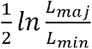, where *L*_*maj*_ and *L*_*min*_ are respectively major and minor axes of the fitted ellipse in Fourier space, and a set of unit vectors ***u*** = (*u*_*x*_,*u*_*z*_) characterizing the anisotropy of pMLC signal on the projected image. Here, *x* and *z* denote cartesian coordinates on the projected image, with *x* oriented along *ϕ* and *z* oriented along 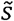 the arclength along the contour of the shape, assuming that the size of the projected window is small compared to the shape radii of curvature. Each unit vector ***u*** is obtained from one interrogation box and is represented by a colored bar in panels such as Fig. 2c (see (*27*) for further details). We computed the associated nematic tensor 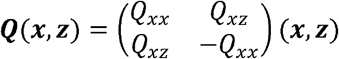, with 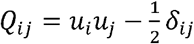 for *i,j* ∈ {*x,z*} and 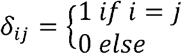. We then averaged row by row the circumferential components *q*(*x*) ≃ ⟨*Q*_*zz*_(*x,z*)⟩*x*, where ⟨ · ⟩*x* denotes averaging along *x*, to obtain profiles of order as a function of *z*. Alternatively, we evaluated a second nematic tensor 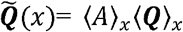, which takes into account both local alignment *A* and average order along *x*, ⟨ *Q* ⟩_*x*_. The cartesian coordinate on projected image *z* is converted to the normalized arclength coordinate on the curved surface 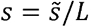, with *L* the length of the shape cross-section, using profiles of average basal endoderm of apical ectoderm surface for each species (Fig. S3C).

#### HCR intensity

HCR intensity profiles *I*(*s*) (Fig. 5C, D, Fig. S9A) were recorded from mid-plane sections using Fiji’s segmented line tool along thick curves inside the ectoderm from the aboral (*s* = 0) to the oral (*s* = 1) pole. These profiles *I*(*s*) were smoothed using a moving average with a window size equivalent to 5% of the curve length, and normalized to set the maximum (respectively, minimum) intensity recorded along the oral-aboral axis at 1 (respectively, 0) and obtain a relative intensity 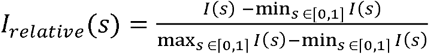.

### “Thompsonian” grids

To highlight the differences between different cnidarian larval shapes, we associate to each shape a deformed grid inspired by D’Arcy Thompson’s famous drawings in (*54*). Those grids are obtained by applying a transformation of the type

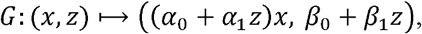

to a common reference grid, *x* and *z* are cartesian coordinates (with *z* the coordinate going along the symmetry axis of the shape cross-section), and *α*_0_, *α*_1_, *β*_0_ and *β*_1_ the parameters of the transformation. This transformation preserves the *x* → − *x* symmetry and the straightness of horizontal and vertical lines (but not the straightness of oblique lines).

For a given axisymmetric shape (*x*(*s*),*z*(*s*)) (with lengths made dimensionless using /?0. the characteristic radius of the shape as unit of length, see SI), we define as a reference shape (*x*_0_(*s*),z_0_(*s*)) corresponding to the unit sphere (for closed shapes) or to a unit spherical cap of same opening radius than the studied shape (for shapes with an oral opening). We then adjust the parameters *α*_0_, *α*_1_, *β*_0_ and *β*_1_ to minimize the objective function 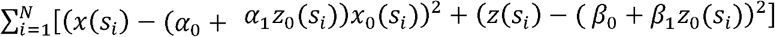 with *s*_*i*_ = *i*/*N* and *N* = 100 a number of sampling points.

We then define a reference grid as a 6-by-6 regular grid circumscribed to the unit circle for closed shapes, and a regular grid circumscribed to a spherical cap with oral opening angle equal to 0.8*π* for open shapes. The grid obtained by applying the associated deformation *G* to the reference grid defines the associated “Thompsonian” grid of the shape.

**Table S1.** Time windows (hours post fertilization) for animal fixation.

|  | Embryo/Gastrula | Early planula | Planula |
| --- | --- | --- | --- |
| *Nematostella*<br>12ppt ASW, 27°C | 18-24 | 30-56 | 66-80 |
| *Aiptasia*<br>36ppt ASW, 27°C | 18-24 | 30-56 | 66-80 |
| *Galaxea*<br>34ppt ASW, 26°C | 33 | 48 | 72 |
| *Acropora*<br>35ppt ASW, 27°C | 28-42 | 54-66 | 78-102 |
| ***Hydractinia***<br>29ppt ASW, 21°C | 10-16 | 18-32 | 48-72 |
| ***Cladonema***<br>28ppt ASW, 20°C | 10-15 | 19-24 | 48-72 |
| ***Nematostella* JNK-in-8**<br>12ppt ASW, 20μM JNK-in-8,<br>0.1% DMSO, 21°C | 22-30 | 44-54 | 66-78 |
| ***Nematostella***<br>**myosinDN**<br>12ppt ASW, 1μM<br>mifepristone, 27°C |  |  | 66-80 |

**Table S2.** Fixation, permeabilization and dehydration methods.

|  | **Fixation**<br>(temperature/incubation time) | **Permeabilization**<br>**/ Wash** | **Dehydration**<br>(serial concentrations) |
| --- | --- | --- | --- |
| ***Nematostella*** | PFA 4%<br>(RT, 1hour) | PTx | Methanol<br>(30-60-100%) |
| ***Aiptasia*** | PFA 4%<br>(RT, 1hour) | PTx | Methanol<br>(30-60-100%) |
| ***Galaxea*** | PFA 4%<br>(RT, 1hour) | PTw | Ethanol<br>(25-50-75-100%) |
| ***Acropora*** | Glutaraldehyde 4%<br>(4°C, 48hours) | PBS | Ethanol<br>(30-50-70-80-90-100%) |
| ***Hydractinia*** | PFA 4%<br>(RT, 1hour) | PTx | Methanol<br>(30-60-100%) |
| ***Cladonema*** | PFA 4%<br>(RT, 1hour) | PBS | Ethanol<br>(25-50-75-100%) |

**Table S3.**
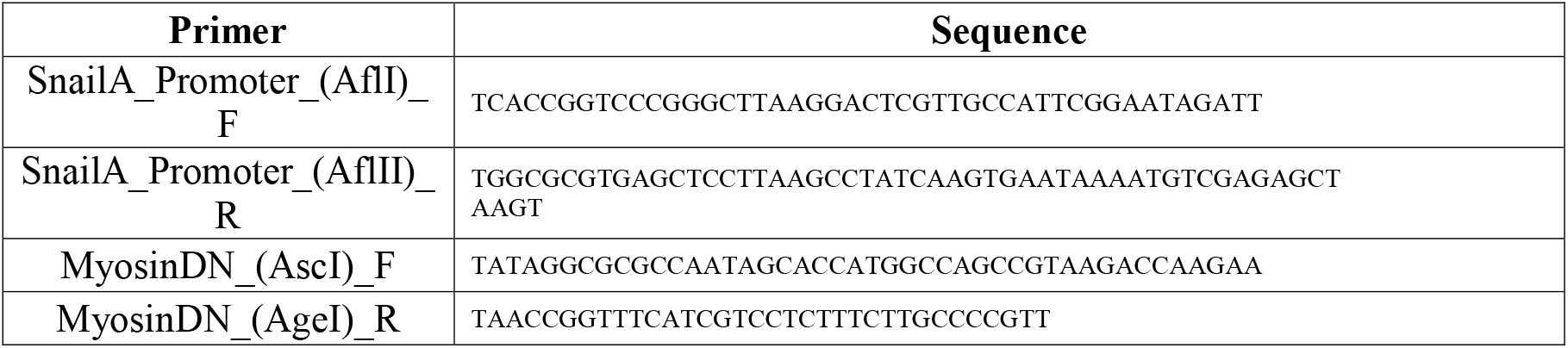
Primers.

**Table S4.** Objectives used per species.

|  | **Objective used for imaging** |
| --- | --- |
| ***Nematostella*** | 40X water (HC PL APO 40X/1.10 W CORR CS2) |
| ***Aiptasia*** | 63X glycerol (HC PL APO 63X/1.30 GLYC CORR CS2)<br>or oil (HC PL APO 63X/1.40 OIL CS2) |
| ***Galaxea*** | 20X air and 63X water |
| ***Acropora*** | 20X air and 63X water |
| *Hydractinia* | 40X water (HC PL APO 40X/1.10 W CORR CS2) |
| *Cladonema* | 63X glycerol (HC PL APO 63X/1.30 GLYC CORR CS2)<br>or oil (HC PL APO 63X/1.40 OIL CS2) |

**Table S5.** Species-specific parameters for image analysis.

| | Distance from peak<br>for normalization of pMLC intensity,<br>D ($\mu\text{m}$) | Size of interrogation<br>boxes<br>for anisotropy analysis,<br>B ($\mu\text{m}$) |
| --- | --- | --- |
| *Nematostella* | 4 | 25 |
| *Aiptasia* | 2 | 10 |
| *Galaxea* | 8 | 40 |
| *Hydractinia* | 4 | 20 |
| *Cladonema* | 2 | 10 |

**Table S6.**
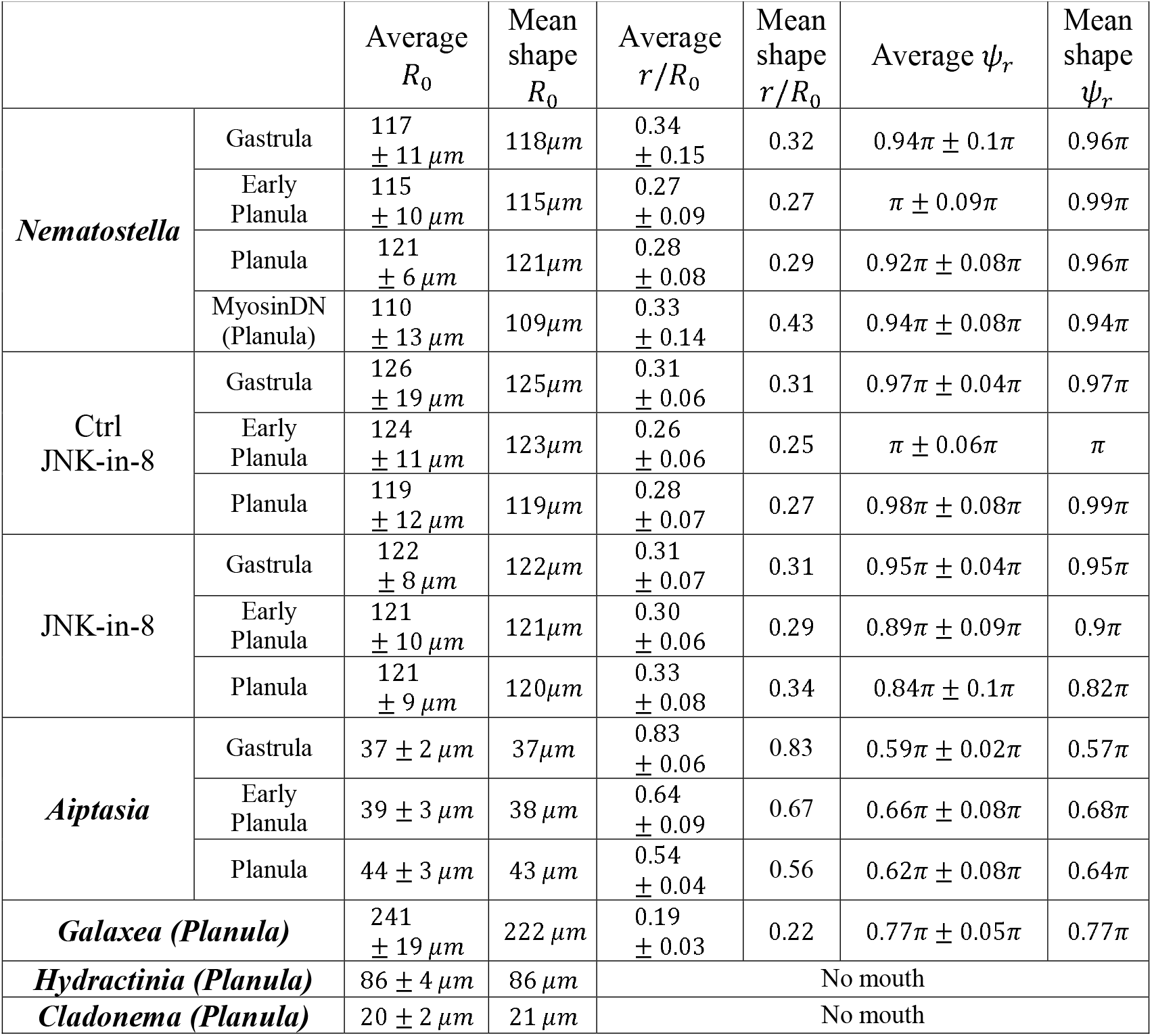
Measured oral parameters for species with oral opening.

| | | Average<br>$R_0$ | Mean<br>shape<br>$R_0$ | Average<br>$r/R_0$ | Mean<br>shape<br>$r/R_0$ | Average $\psi_r$ | Mean<br>shape<br>$\psi_r$ |
| --- | --- | --- | --- | --- | --- | --- | --- |
| *Nematostella* | Gastrula | 117<br>$\pm 11 \mu\text{m}$ | 118$\mu\text{m}$ | 0.34<br>$\pm 0.15$ | 0.32 | $0.94\pi \pm 0.1\pi$ | 0.96π |
| | Early Planula | 115<br>$\pm 10 \mu\text{m}$ | 115$\mu\text{m}$ | 0.27<br>$\pm 0.09$ | 0.27 | $\pi \pm 0.09\pi$ | 0.99π |
| | Planula | 121<br>$\pm 6 \mu\text{m}$ | 121$\mu\text{m}$ | 0.28<br>$\pm 0.08$ | 0.29 | $0.92\pi \pm 0.08\pi$ | 0.96π |
| | MyosinDN<br>(Planula) | 110<br>$\pm 13 \mu\text{m}$ | 109$\mu\text{m}$ | 0.33<br>$\pm 0.14$ | 0.43 | $0.94\pi \pm 0.08\pi$ | 0.94π |
| Ctrl<br>JNK-in-8 | Gastrula | 126<br>$\pm 19 \mu\text{m}$ | 125$\mu\text{m}$ | 0.31<br>$\pm 0.06$ | 0.31 | $0.97\pi \pm 0.04\pi$ | 0.97π |
| | Early Planula | 124<br>$\pm 11 \mu\text{m}$ | 123$\mu\text{m}$ | 0.26<br>$\pm 0.06$ | 0.25 | $\pi \pm 0.06\pi$ | $\pi$ |
| | Planula | 119<br>$\pm 12 \mu\text{m}$ | 119$\mu\text{m}$ | 0.28<br>$\pm 0.07$ | 0.27 | $0.98\pi \pm 0.08\pi$ | 0.99π |
| JNK-in-8 | Gastrula | 122<br>$\pm 8 \mu\text{m}$ | 122$\mu\text{m}$ | 0.31<br>$\pm 0.07$ | 0.31 | $0.95\pi \pm 0.04\pi$ | 0.95π |
| | Early Planula | 121<br>$\pm 10 \mu\text{m}$ | 121$\mu\text{m}$ | 0.30<br>$\pm 0.06$ | 0.29 | $0.89\pi \pm 0.09\pi$ | 0.9π |
| | Planula | 121<br>$\pm 9 \mu\text{m}$ | 120$\mu\text{m}$ | 0.33<br>$\pm 0.08$ | 0.34 | $0.84\pi \pm 0.1\pi$ | 0.82π |
| *Aiptasia* | Gastrula | 37 $\pm 2 \mu\text{m}$ | 37$\mu\text{m}$ | 0.83<br>$\pm 0.06$ | 0.83 | $0.59\pi \pm 0.02\pi$ | 0.57π |
| | Early Planula | 39 $\pm 3 \mu\text{m}$ | 38 $\mu\text{m}$ | 0.64<br>$\pm 0.09$ | 0.67 | $0.66\pi \pm 0.08\pi$ | 0.68π |
| | Planula | 44 $\pm 3 \mu\text{m}$ | 43 $\mu\text{m}$ | 0.54<br>$\pm 0.04$ | 0.56 | $0.62\pi \pm 0.08\pi$ | 0.64π |
| *Galaxea* (Planula) | | 241<br>$\pm 19 \mu\text{m}$ | 222 $\mu\text{m}$ | 0.19<br>$\pm 0.03$ | 0.22 | $0.77\pi \pm 0.05\pi$ | 0.77π |
| *Hydractinia* (Planula) | | 86 $\pm 4 \mu\text{m}$ | 86 $\mu\text{m}$ | No mouth | | | |
| *Cladonema* (Planula) | | 20 $\pm 2 \mu\text{m}$ | 21 $\mu\text{m}$ | No mouth | | | |

