## Supplemental Figures for "Cross-Species Biomechanical Determinants of Shape Diversity"

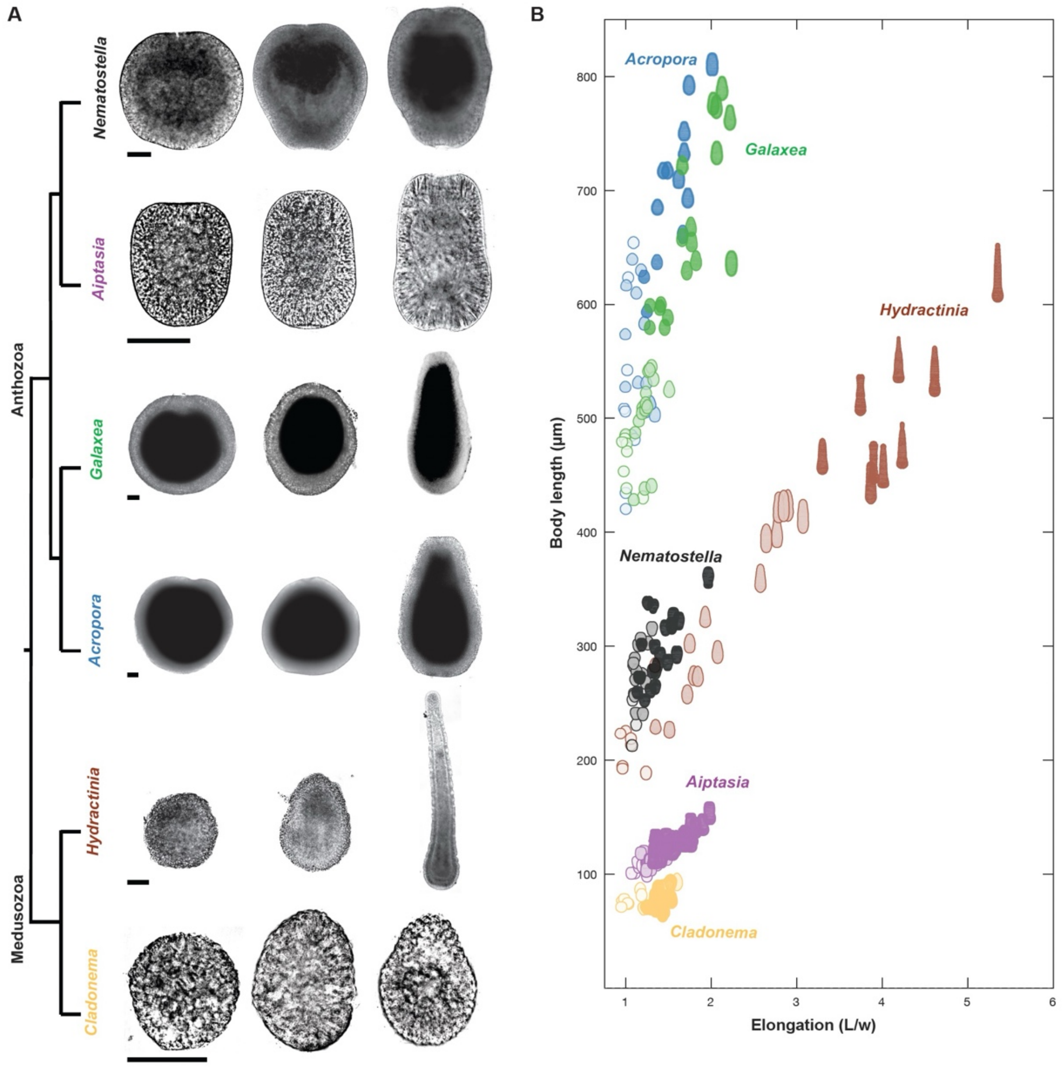


**Fig. S1. Diversity of cnidarian shapes. (A)** Simplified phylogenetic tree of Cnidaria with representative bright field images selected over three developmental stages (see Table S1). Scale bars: 50µm. **(B)** Morphospace of cnidarian larval shapes, positioned by their degree of elongation and body length**.** Segmented shapes are obtained from bright field images.

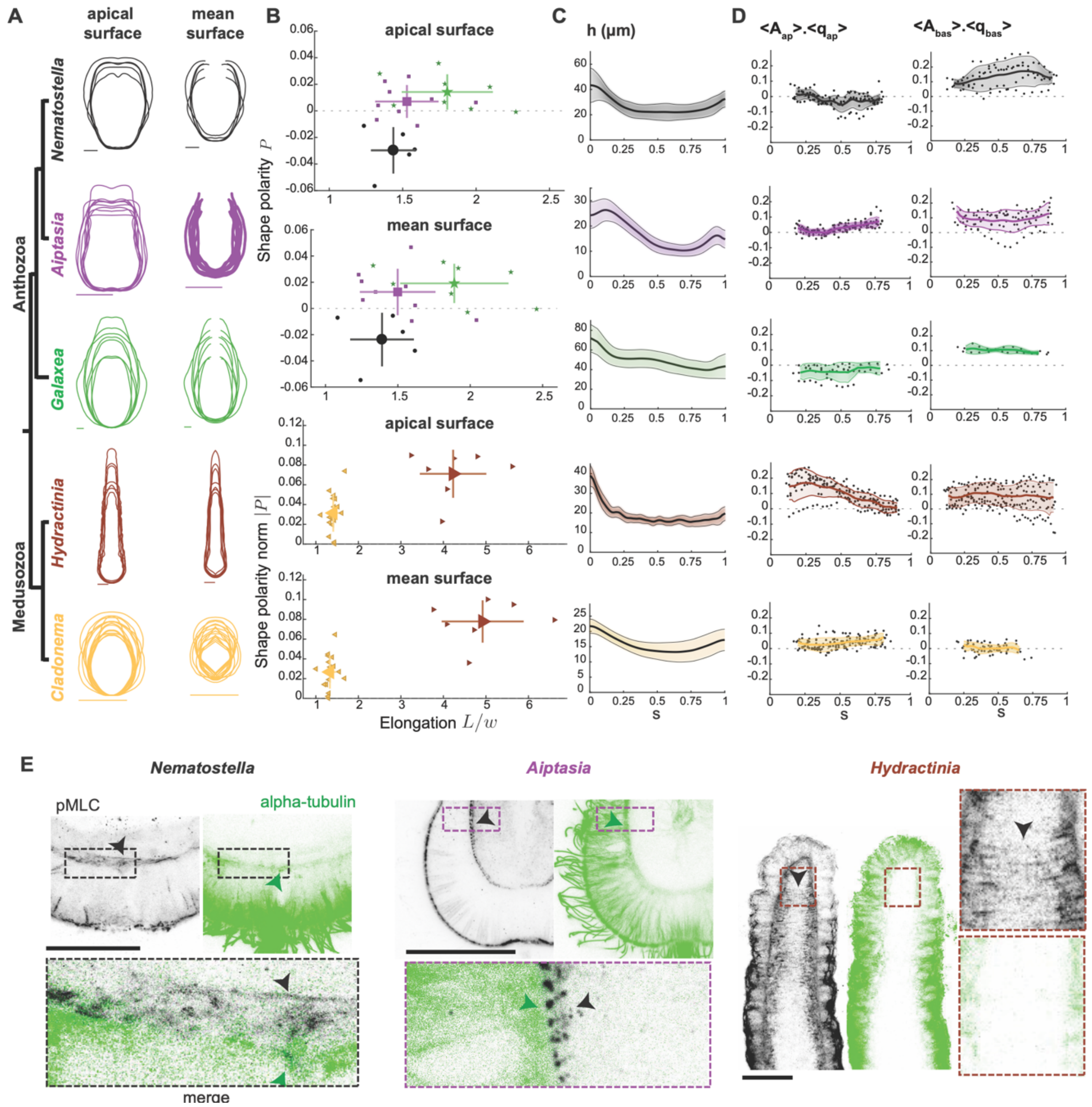


**Fig. S2. Ectodermal thickness and shapes of apical, basal and mean surfaces in cnidarians.** (**A**) Individual symmetrized curves extracted from cross-sections images of pMLC immunostainings overlapped for the apical surface (Left) and mean surface (Right) at the planula stage for the apical (left) and mean (right) surfaces (see Material and Methods). **(B)** Shape polarity and elongation of apical (left) and mean (right) surfaces for anthozoans (top) and for medusozoans (bottom, absolute value of shape polarity is used). Shapes are obtained from pMLC immunostainings at planula stage. All scale bars represent 50µm. **(C)** Profiles of ectodermal thickness as a function of normalized arclength coordinate, averaged over the planula stage. **(D)** Quantifications of circumferential order as the product of local alignment, $A$, and nematic order parameter, $q$, in the apical (left) and basal (right) surfaces as a function of normalized arclength coordinate, averaged over the planula stage. Shaded areas in C and D indicate standard deviations. **(E)** Double staining of alpha-tubulin and pMLC in *Nematostella*, *Aiptasia* and *Hydractinia* planulae. Alpha-tubulin is enriched in the ectoderm (green arrowheads) while pMLC is enriched at the basal interface between the ectoderm and the endoderm (black arrowheads).

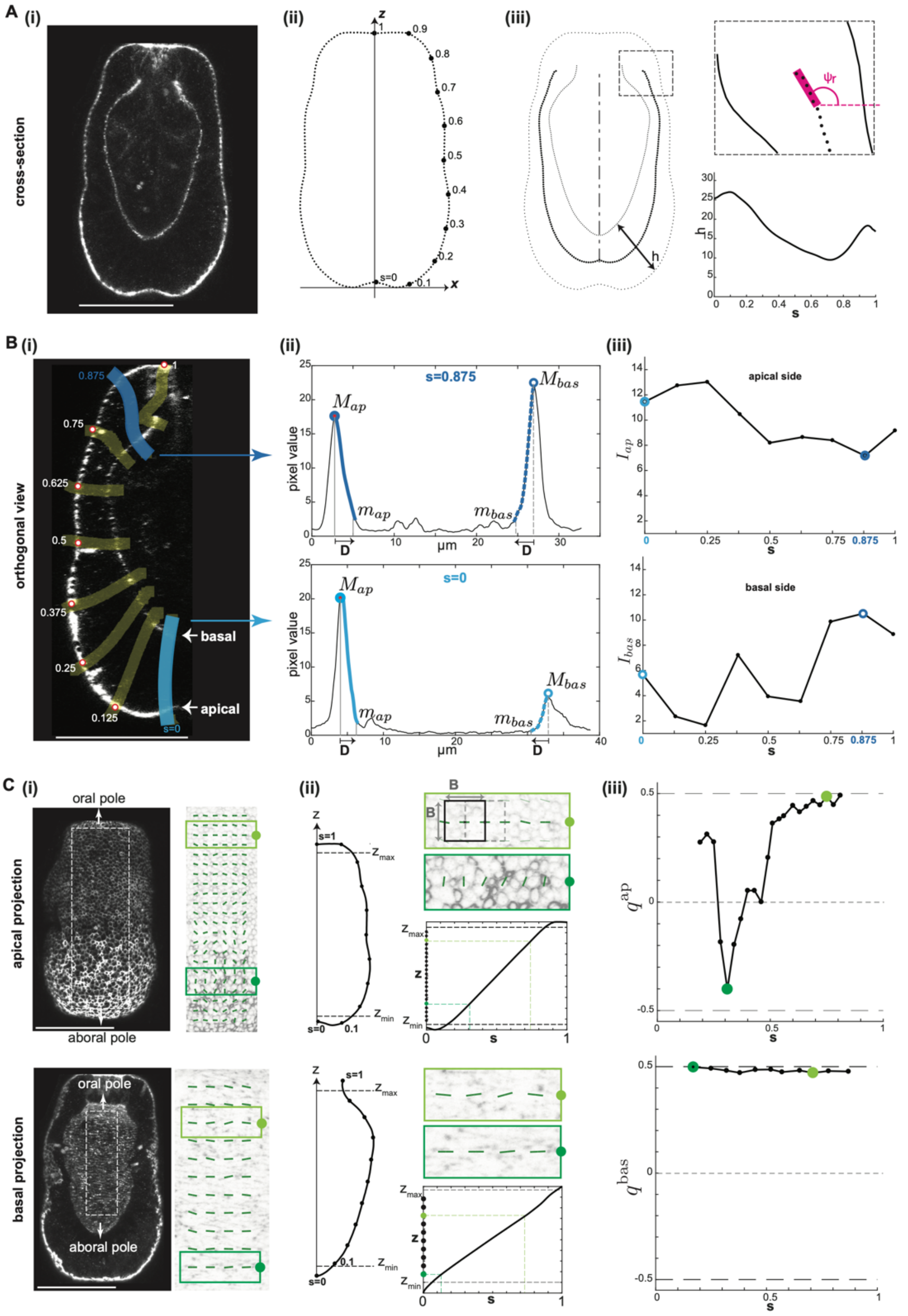


**Fig. S3. Segmentation and quantification. (A)** pMLC cross-sections images **(i)** were used to delineate the apical and basal surfaces, define the normalized arclength coordinate $s$ **(ii)**, fit the oral boundary angle $\psi_r$, and compute thickness h **(iii)**. **(B) (i)** Orthogonal view with overlapping *s* positions (red dots) where pMLC intensity was probed using Fiji's Segmented Line tool (thick yellow lines; thick blue lines are detailed as examples). **(ii)** Examples of pixel intensity curves along the apico-basal axis, drawn at positions s=0 (bottom) and s=0.875 (top) showing the local maxima, $M_{ap}$ and $M_{bas}$, corresponding to the signal-enriched apical and basal surfaces, and the domains used for normalization by $m_{ap}$ and $m_{bas}$ at a distance $D$ of the peaks (see material and methods and Table S5). **(iii)** Plots showing the resulting normalized pMLC intensity as a function of *s* for the apical ($I_{ap}$, Top) and basal ($I_{bas}$, Bottom) surfaces. (**C) (i)** pMLC apical (top) and basal (bottom) projections with dashed regions used to compute coarse-grained anisotropy. **(ii)** Coordinate mapping for apical (top) and basal (bottom) surfaces. For each case: (Left) half-shapes with *z*-axis and *s* coordinate landmarks. (Top right) Example of two adjacent interrogation windows and box size parameter B

(Table S5). (Bottom right) Function $z(s)$ used for inverse coordinate mapping from $z$ to $s$, with dark and pale green lines corresponding to centers of boxes shown on top. **(iii)** Resulting nematic order as a function of $s$ for the apical ($q^{ap}$, top) and basal ($q^{bas}$, bottom) surfaces. Scale bars: 50μm.

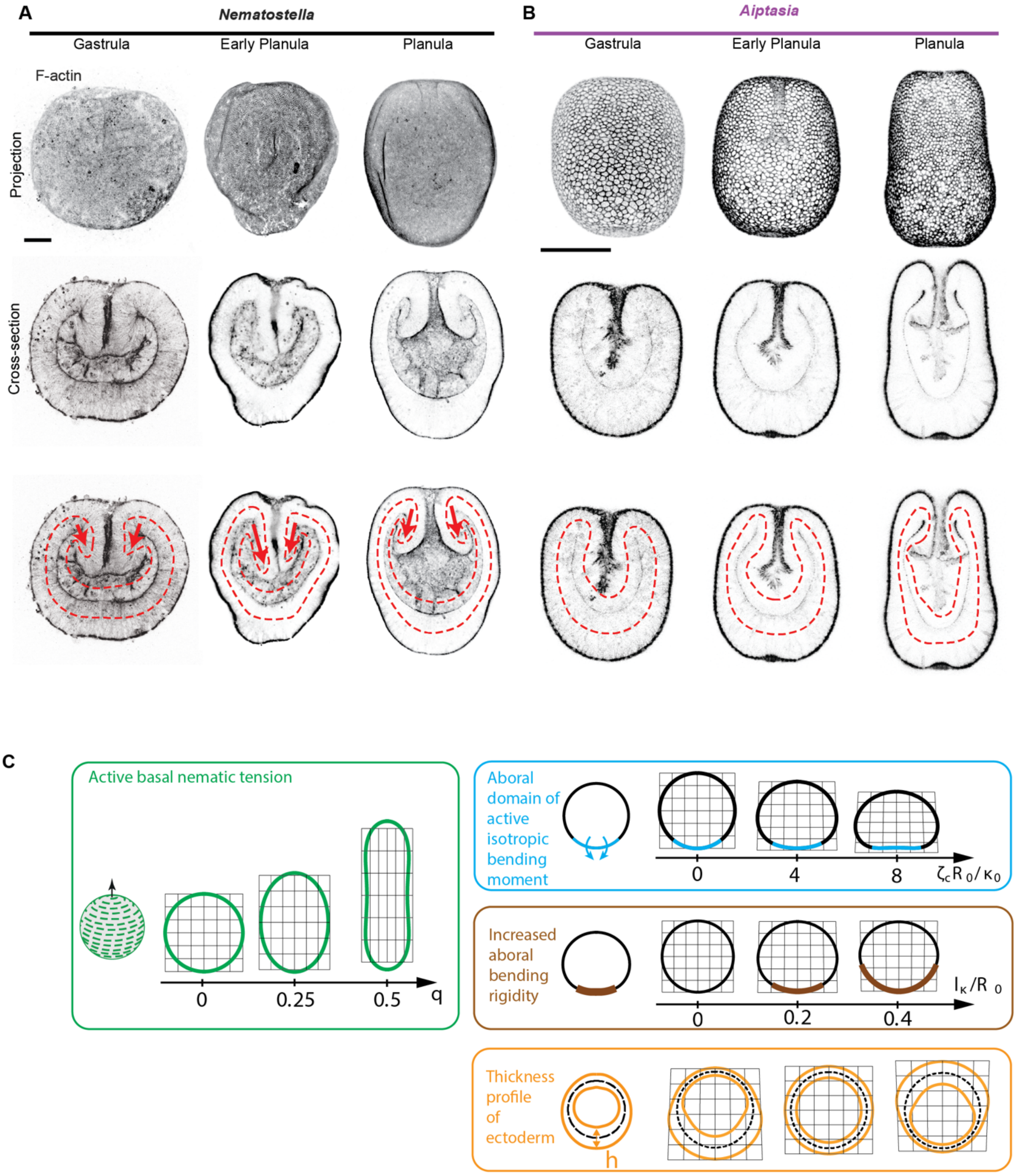


**Fig. S4. *Nematostella* and *Aiptasia* oral geometry, and biomechanical modules in closed shells (A), (B).** Cross-sections and projections of phalloidin staining (labelling F-actin) in three developmental stages of *Nematostella* (A) and *Aiptasia* (B). Red lines roughly indicate mean surfaces of ectoderm and endoderm. Arrows indicate oral tissue involution. (**C**) Biomechanical modules underlying tissue elongation and shape polarity; as in Fig. 3B-F, but for closed surfaces. Shapes correspond to steady-state simulations of the active surface model. Modules correspond to variations in basal nematic order aboral isotropic bending moment, increased aboral bending rigidity and ectoderm thickness. Scale bars: 50µm.

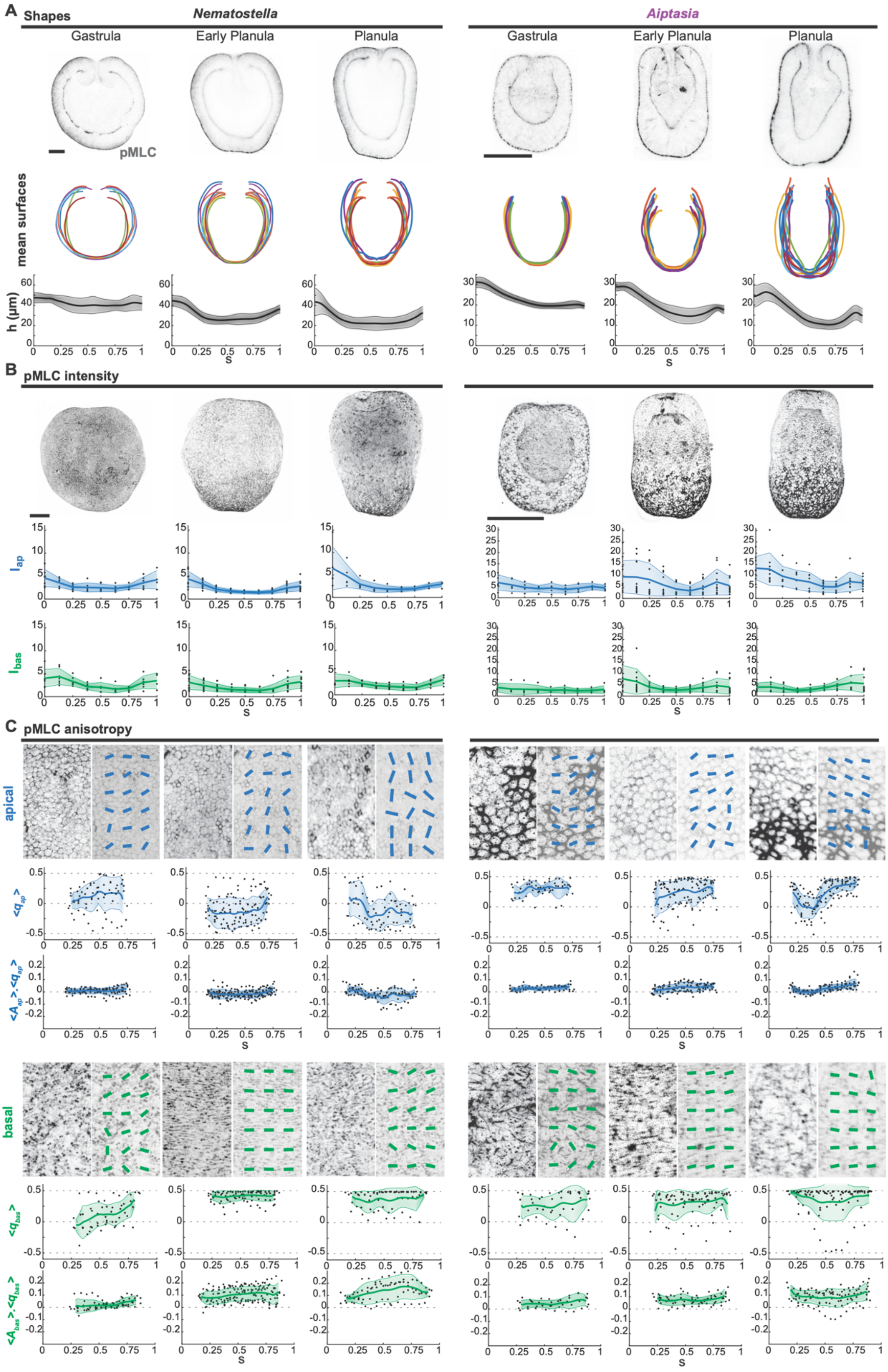


**Fig. S5. Myosin intensity and anisotropy during larval development of *Nematostella* and *Aiptasia.* (A)** (Top) Images of gastrula, early planulae and planulae stained for pMLC in *Nematostella* (Left) and *Aiptasia* (Right). (Middle) Computed mean surfaces overlapped for each stage. (Bottom) Quantification of ectodermal thickness averaged over each stage, as a function of normalized arclength coordinate *s*. **(B)** (Top) Max projections of pMLC immunostainings for the three stages. (Bottom) pMLC intensity profile within the apical and basal surfaces as a function of normalized arclength coordinate $s$, over the three stages. **(C)** pMLC anisotropy dynamics. (Top) Zoom-in views of apical and basal surfaces with superimposed bars representing anisotropy orientation (Material and Methods). (Bottom) Quantification of circumferential order in the apical and basal surfaces as a function of normalized arclength coordinate. Circumferential order is quantified according to the nematic order parameter $q$ or by the product of local alignment $A$ and nematic order $q$ (Material and Methods). Shaded areas indicate standard deviations. Scale bars: 50μm.

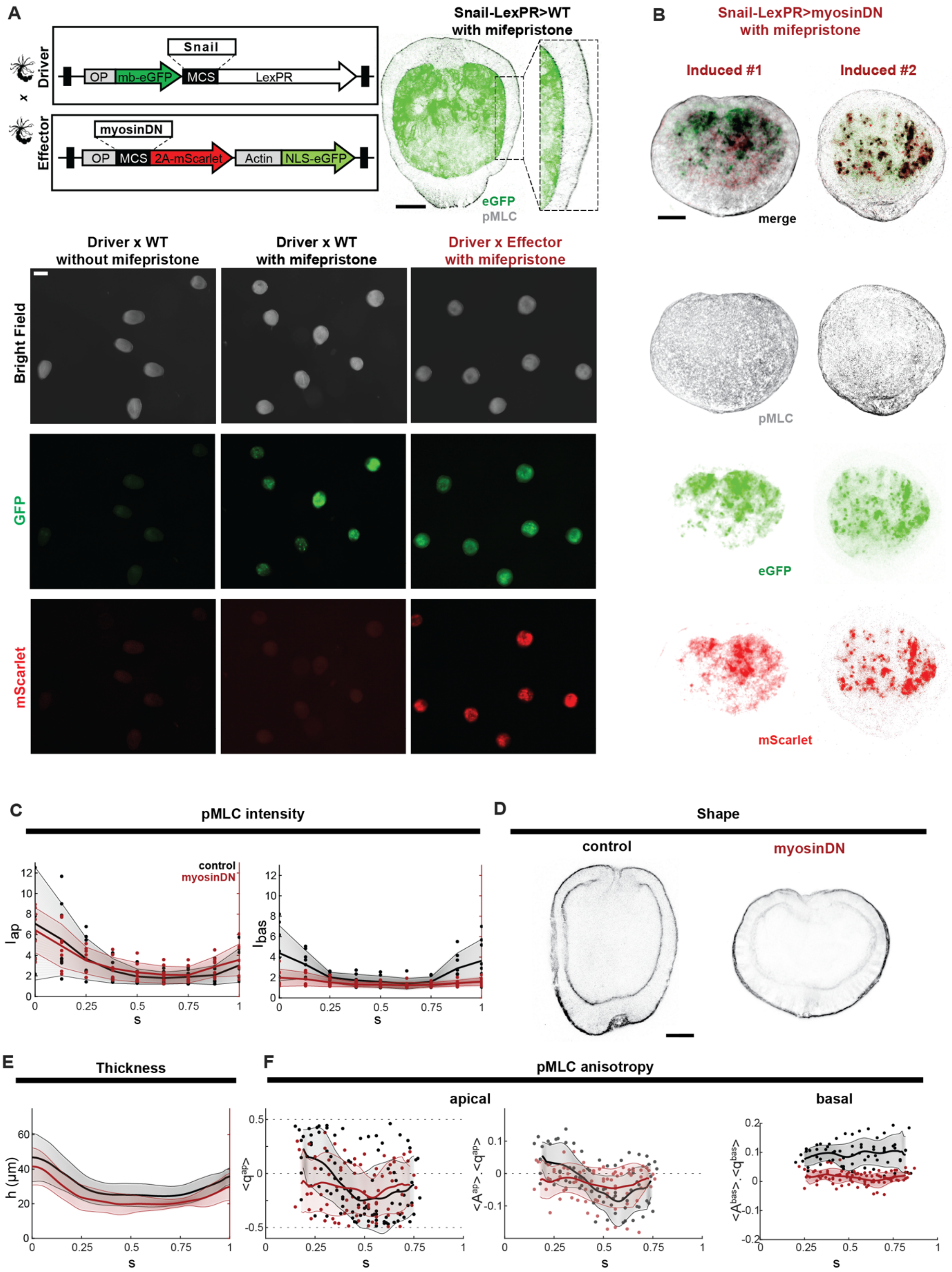


**Fig. S6. Detailed description of inducible genetic system. (A)** Schematic representation of the inducible LexA-based gene expression system. We used a modular expression system utilizing separate driver and effector cassettes. (Top-Right) Confocal image of an induced control animal carrying *LexPR>OP-mb-eGFP*, showing membrane-localized eGFP (mb-eGFP) expression in the endoderm, combined with pMLC staining. The inset displays a zoomed-in view, highlighting that the strong basal pMLC signal does not overlap with the endodermal membrane-localized eGFP. Scale bar: 50μm. (Bottom) Stereoscope images showing bright field (top row) and no GFP (middle row) and mScarlet (bottom row) expression in Snail-LexPR>WT planula with no mifepristone in the medium (left column), GFP expression in mifepristone treated Snail-LexPR>WT planula (middle column), and both GFP and mScarlet in mifepristone treated Snail-LexPR>OP-myosinDN-2A-mScarlet induced planula (right column). Scale bar: 200μm. Mifepristone treated Snail-LexPR>WT animals elongate, while mifepristone treated Snail-LexPR>OP-myosinDN-2A-mScarlet animals fail to elongate (**B**) Projection images of two Snail-LexPR>OP-myosinDN-2A-mScarlet induced planulae immuno-stained against pMLC, GFP, and mScarlet antibodies. Scale bar: 50μm**. (C)** Profiles of pMLC intensity as a function of normalized arclength coordinate within the apical surface in control (black) and Snail-LexPR>OP-myosinDN induced (red) planulae. **(D)** Cross-section image of a control (left) and a Snail-LexPR>OP-myosinDN induced (right) planula stained with pMLC. **(E)** Profiles of ectodermal thickness as a function of normalised arclength coordinate $s$, averaged over the planula stage in control (black) and Snail-LexPR>OP-myosinDN induced (red) planula. **(F)** pMLC anisotropy in Snail-LexPR>OP-myosinDN induced *Nematostella*. (Top-left) Zoom-in view of the apical surface with superimposed bars representing anisotropy orientation in control and Snail-LexPR>OP-myosinDN induced planulae. (Bottom-left) Profiles of nematic order parameter $q$ and product of local alignment $A$ and nematic order $q$ in the apical surface as a function of $s$, in control (black) and induced (red) animals. (Right) Profiles of product of local alignment $A$ and nematic order $q$ in in the basal surface as a function of normalized arclength coordinate $s$ in control (black) and induced (red) animals. Shaded areas indicate standard deviation.

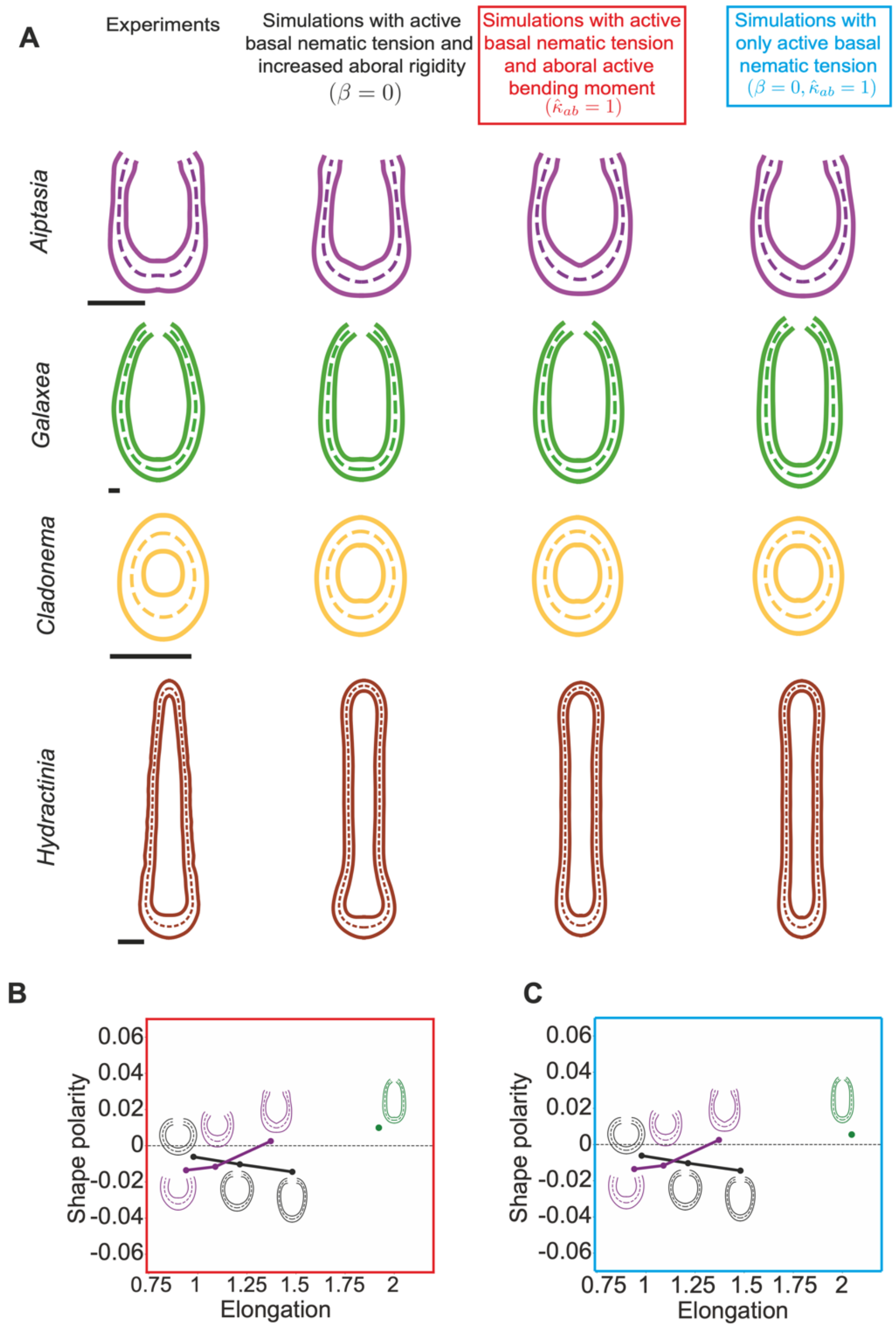


**Fig. S7. Simulations of larval shapes with active basal nematic tension, and with and without increased aboral rigidity and aboral active bending moment. (A)** Experimental shapes and fitted simulated shapes with active basal nematic tension and increased aboral rigidity, active basal nematic tension and aboral active bending moment, and active basal nematic tension alone. See SI for additional details and SI Table 5 for simulation parameters. Scale bars: 50μm. **(B),(C)** Morphospace of simulated larval optimal shapes of *Nematostella*, *Aiptasia* and *Galaxea,* with active basal nematic tension and aboral active bending moment (B) and active basal nematic tension alone (C). Shapes are positioned according to outer surface elongation and shape polarity, as in Fig. 5I.

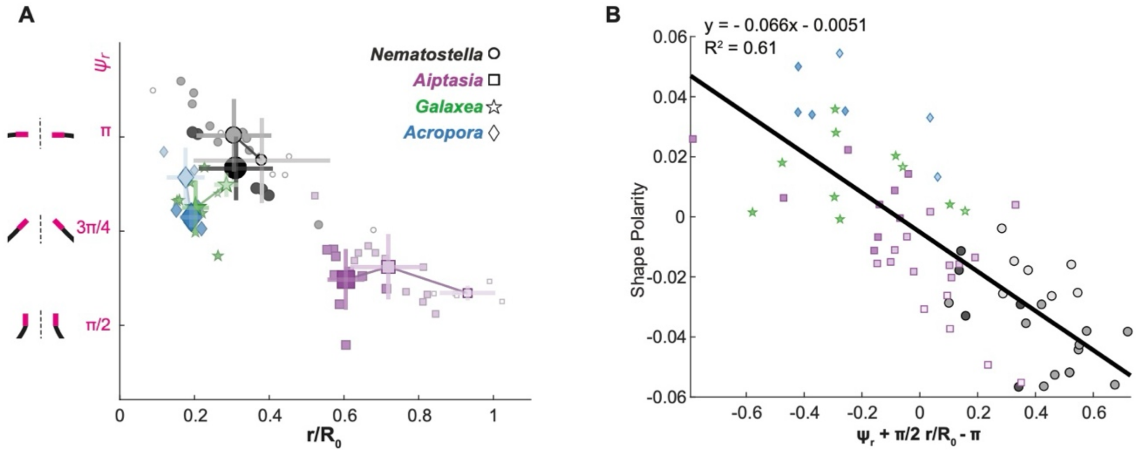


**Fig. S8. Dynamics of oral opening geometry, and relationship to shape polarity.** (**A**). Dynamics of oral opening geometry (measured by the angle $\psi_r$ and normalized opening radius $r/R_0$) over developmental stages for *Nematostella*, *Aiptasia*, *Galaxea*, *Acropora*, with color code indicated on the right. Darker colors correspond to early larval, larva and planula. (**B**) Correlation between shape polarity and opening angle for *Nematostella*, *Aiptasia*, *Galaxea*, *Acropora*, at planula stage.

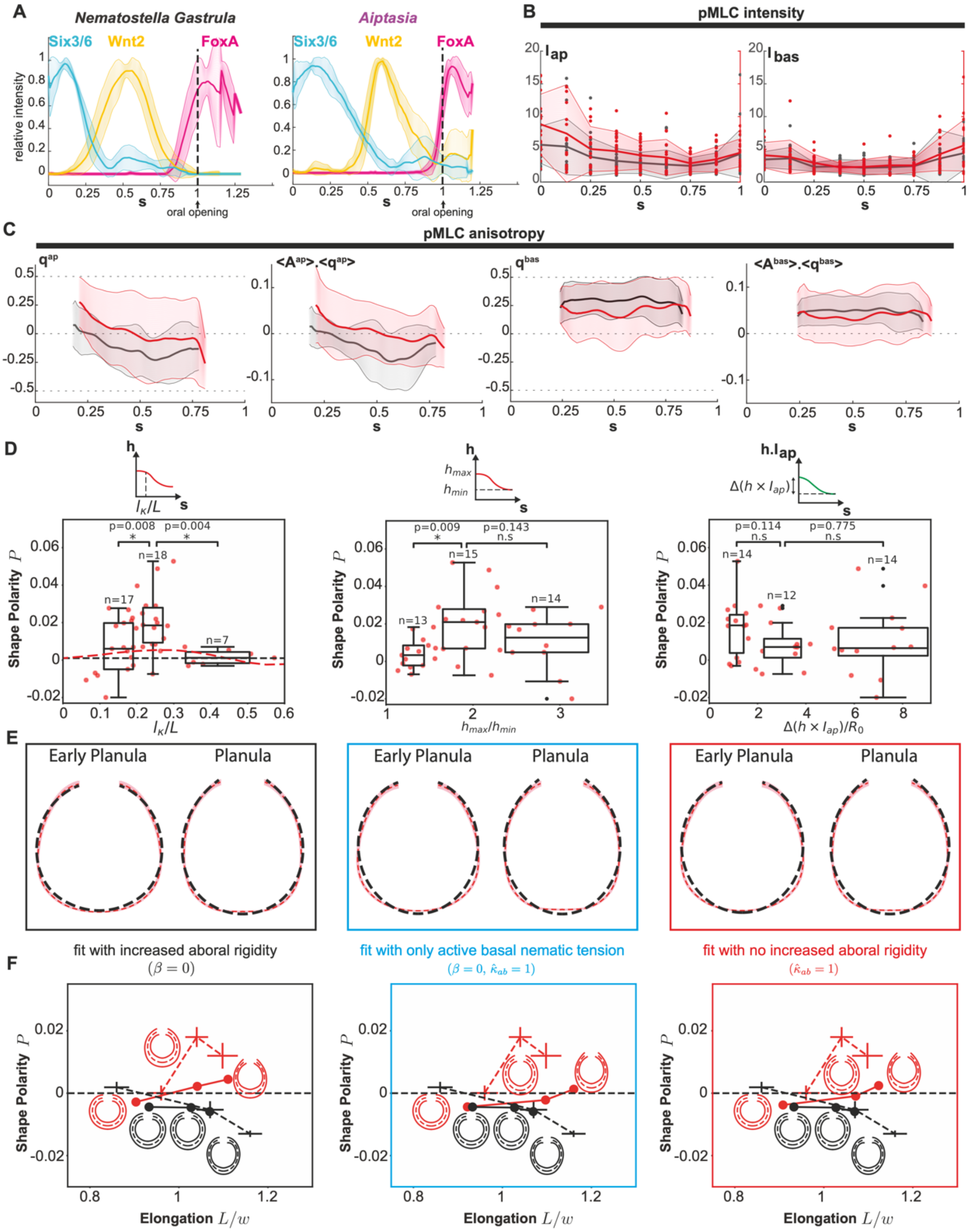


**Fig. S9. Perturbation of axial patterning in *Nematostella*. (A)** Normalized HCR intensity of Six3/6, Wnt2 and FoxA as a function of normalized arclength coordinate $s$, in *Nematostella* and *Aiptasia* gastrulas. **(B-C)** Profiles of pMLC intensity (B), nematic order parameter (C), and product of local alignment $A$ and nematic order $q$ (C) as a function of normalized arclength coordinate $s$, for apical and basal surfaces, in a set of *Nematostella* planulae treated with JNK-in-8 (red) and corresponding control planulae (black). (**D**) Mean shell polarity as a function of the size of the thicker aboral region (left), the ratio between maximum and minimum ectoderm thickness (center) and the difference between the maximum and the minimum of the product between ectoderm thickness and normalized apical pMLC intensity (right) in a set of *Nematostella* treated with JNK-in-8 at all stages (Gastrula, Early Planula and Planula). Red dots: data for individual larvae. Box plots: Binned dataset. The box extends vertically from the first quartile to the third quartile of the polarity distribution of the sample, with a line at the median. The whiskers extend from the box to the farthest data point lying within 1.5 times the inter-quartile range from the box. Additional black points are data points beyond the end of the whiskers. Each boxplot is centered around the horizontal mean of the corresponding sample, with horizontal width equal to two times the horizontal standard deviation of the sample. Means of each sample are compared by performing a two-sided t-test. The difference between the means is significant (indicated by *) if the p-value of the test is less than 5% and non-significant (indicated by n.s) otherwise. Red dotted curve in left graph: mean shell polarity as a function of the size of the aboral increased rigidity domain in simulations, with oral geometry and nematic basal tension parameters as for JNK Planula (Figure 5G). Black dotted line: mean shell polarity for the same parameters as the red line, but in the absence of increased aboral rigidity. (**E**) Mean experimental shapes (dotted red, pink shadow represent the standard error on the mean of the experimental shapes) and optimal simulated shapes (dotted black) of *Nematostella* treated with JNK-in-8 at Early Planula and Planula stages. Optimal simulated shapes are obtained as follows. Black box (left): fit on $\hat{\kappa}_{ab}$ with the values of $\beta$ and $\beta_n$ obtained by fitting the data on *Nematostella* wild type and control. Blue box (center): fit on $\beta_n$, imposing $\beta = 0$ and $\hat{\kappa}_{ab} = 1$. Red box (right): fit on $\beta$, imposing $\hat{\kappa}_{ab} = 1$ and with the value of $\beta_n$ obtained by fitting the data on *Nematostella* wild type and Control (right). See SI section 3.5 for details. (**F**) Morphospace showing larval shapes of developing *Nematostella* (control, dotted black; JNK-in-8 treated, dotted red) and corresponding optimal simulations (control, black; JNK-in-8 treated, red) positioned according to elongation and shape polarity. Error bars on experimental data represent the standard error of the mean. Results for the corresponding fits in (E) are displayed with colored boxes indicating the correspondence.

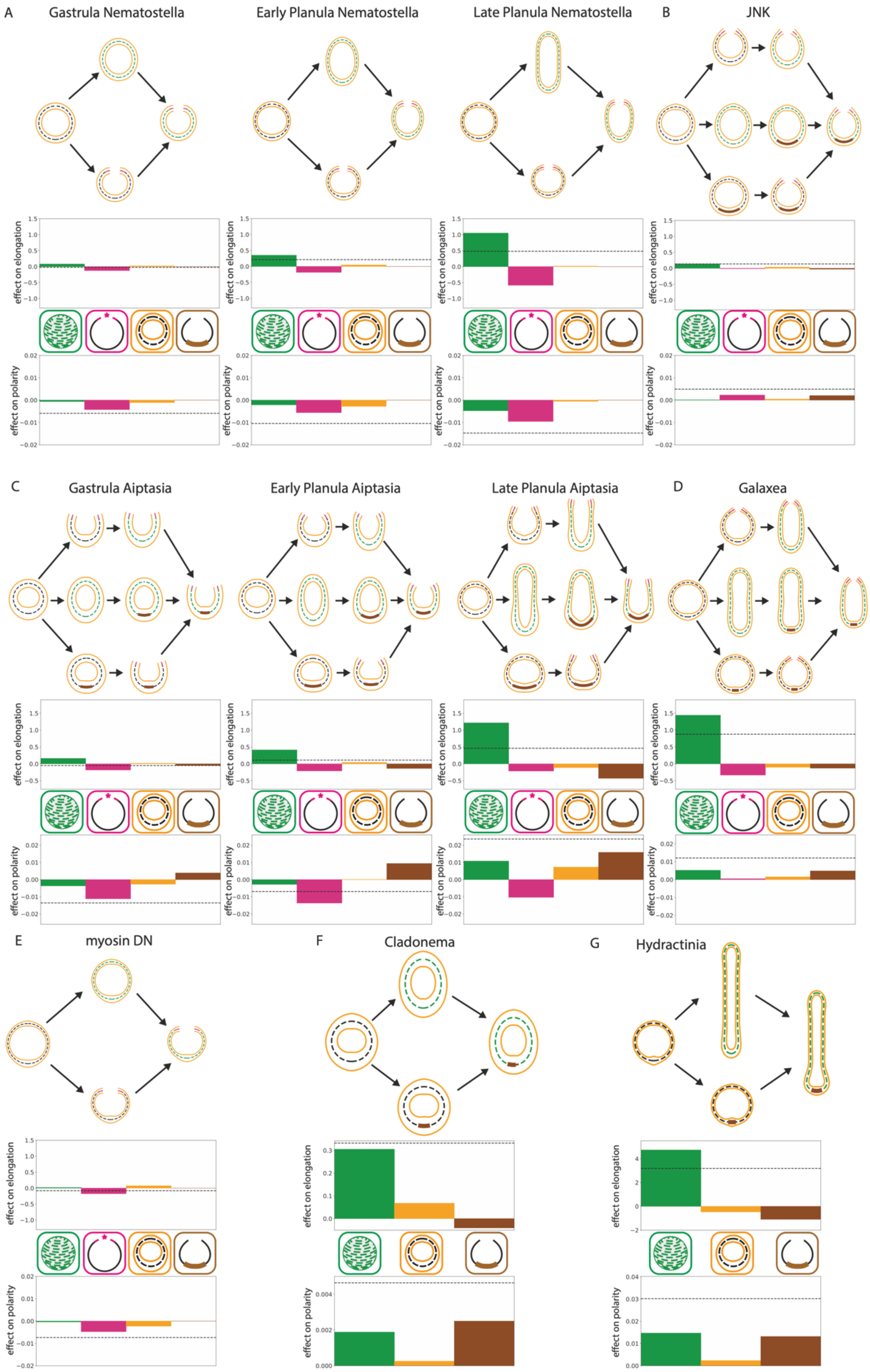


**Fig. S10. Contributions of biomechanical modules in active surface simulations of cnidarian shapes.** For each panel: (top) Simulated shapes where two or three modules (oral opening, basal nematic, aboral rigidity) are added sequentially in different orders; (bottom) relative average contribution of four modules (species with open mouth) or three modules (species with closed mouth) on elongation and shape polarity. For simplicity, the effect of the thickness module is not shown on simulated shapes on top. **(A)** *Nematostella* at gastrula, early planula and late planula stage, **(B)** JNK-treated *Nematostella* at late planula stage, **(C)** *Aiptasia* at gastrula, early planula and late planula stage, **(D)** *Galaxea* at late planula stage, **(E)** myosin-DN *Nematostella* at late planula stage, **(F)** *Cladonema* at late planula stage and **(G)** *Hydractinia* at late planula stage. See SI for further details.

### Movie captions

**Movie S1**: Simulated steady-state shape as a function of nematic order $q$, for an active basal nematic tension profile acting on a spherical cap with opening angle $\psi_r$ =0.8$\pi$. The nematic order is uniform away from the poles. See SI section 3.2 for other model parameters.

**Movie S2**: Simulated steady-state shape of a passive open shell as a function of the opening angle $\psi_r$. The opening radius $r$ is constant and equal to the opening radius of a spherical cap with opening angle $\psi_r$ =0.8$\pi$. See SI section 3.2 for other model parameters.

**Movie S3**: Simulated steady-state shape as a function of the magnitude $\zeta_c$ of the aboral gradient of isotropic active bending moment The profile of isotropic active bending moment is taken piecewise-constant, and vanishes except in the region $0 \leq s < 0.3$ where it is set to $\zeta_c$ .The reference shape for $\zeta_c$=0 is a spherical cap with opening angle $\psi_r$ =0.8$\pi$. See SI section 3.2 for other model parameters.

**Movie S4**: Simulated steady-state shape as a function of the size of the increased aboral rigidity domain $l_\kappa$. The region $0 \leq s < l_\kappa/L$ has increased bending rigidity $\kappa_{ab}/\kappa_0 = 2$. The reference shape with uniform bending rigidity is a spherical cap with opening angle 0.8 $\pi$. See SI section 3.2 for other model parameters.

**Movie S5**: Variation of the outer surface shape with the ectoderm thickness profile. The outer surface is obtained from the thickness profile shown on the right. The mean surface is a spherical cap with opening angle $\psi_r$ =0.8$\pi$. See SI section 3.2 for other model parameters.
