## Supplementary material for "Cross-Species Biomechanical Determinants of Shape Diversity": SI

### Cross-Species Biomechanical Determinants of Shape Diversity: Supplementary Theory

February 7, 2025

#### 1 Active patterned viscoelastic shell model

In this section we introduce an active surface model for the surface mechanics of cnidarian larvae.

##### 1.1 Geometry of the surface

We describe the tissue by an infinitely thin surface that can be associated to the ectoderm mean surface, defined as the mid-surface between the apical surface of the ectoderm and the basal surface of the ectoderm/endoderm. pMLC stainings indicate that myosin is largely expressed in the ectoderm and basal surface of the ectoderm/endoderm (Fig. 2A), so that we assume that the endoderm contribute mechanically only through myosin present at its basal interface.

We consider a cartesian basis $\mathbf{e}_x$, $\mathbf{e}_y$, $\mathbf{e}_z$, and an axisymmetric surface $\mathcal{S}$ with position determined by

$$\boldsymbol{X}(\phi, s) = (x(s)\cos\phi, x(s)\sin\phi, z(s)), \tag{1}$$

where $\phi \in [0, 2\pi]$ is the azimuthal angle and $s \in [0, L]$ the arclength coordinate such that $|\partial_s \boldsymbol{X}| = 1$ (Fig. 3A). At $s = 0$ (aboral pole) the surface is closed, while at $s = L$ (oral pole) the surface can be open or closed depending on the species. We also introduce the angle $\psi$ defined such that $\partial_s x = \cos\psi$ and $\partial_s z = \sin\psi$.

**Fundamental tensors.** We follow notations of differential geometry as in Refs [2, 3]. The local tangent basis $(\boldsymbol{e}_\phi, \boldsymbol{e}_s)$ and the normal vector $\boldsymbol{n}$ are given by:

$$\boldsymbol{e}_\phi = \partial_\phi \boldsymbol{X} = (-x(s)\sin\phi, x(s)\cos\phi, 0), \tag{2}$$

$$\boldsymbol{e}_s = \partial_s \boldsymbol{X} = (\cos\phi\cos\psi, \sin\phi\cos\psi, \sin\psi), \tag{3}$$

$$\boldsymbol{n} = \frac{\boldsymbol{e}_\phi \times \boldsymbol{e}_s}{|\boldsymbol{e}_\phi \times \boldsymbol{e}_s|} = (\cos\phi\sin\psi, \sin\phi\sin\psi, -\cos\psi). \tag{4}$$

The metric, inverse metric, curvature and antisymetric Levi-Civita tensors read

$$g_{ij} = \begin{pmatrix} x^2 & 0 \\ 0 & 1 \end{pmatrix}, \; g^{ij} = \begin{pmatrix} 1/x^2 & 0 \\ 0 & 1 \end{pmatrix}, \; C_i^j = \begin{pmatrix} \frac{\sin\psi}{x} & 0 \\ 0 & \partial_s\psi \end{pmatrix}, \; \epsilon_{ij} = \begin{pmatrix} 0 & x \\ -x & 0 \end{pmatrix}. \tag{5}$$

and we can show the following useful relations:

$$\partial_s C_\phi^\phi = \frac{\cos\psi}{x}\left(C_s^s - C_\phi^\phi\right), \tag{6}$$

$$\partial_s\left(\frac{\cos\psi}{x}\right) = -C_\phi^\phi C_s^s - \left(\frac{\cos\psi}{x}\right)^2. \tag{7}$$

**Christoffel symbols and covariant derivatives.** The Christoffel symbols of second kind are defined as $\Gamma^k_{ij} = \partial_i \boldsymbol{e}_j \cdot \boldsymbol{e}^k$. There are three non-zero Christoffel symbols in our case:

$$\Gamma^s_{\phi\phi} = -x\cos\psi,\ \Gamma^\phi_{s\phi} = \Gamma^\phi_{\phi s} = \frac{\cos\psi}{x}. \tag{8}$$

The covariant derivative of a vector $\mathbf{a}$ is defined as $\nabla_i a^j = \partial_i a^j + \Gamma^j_{ik} a^k$ and the covariant derivative of a tensor $\boldsymbol{A}$ is defined as $\nabla_i A^{jk} = \partial_i A^{jk} + \Gamma^j_{il} A^{lk} + \Gamma^k_{il} A^{jl}$. We assume that all physical quantities are $\phi$-independent, we consider vectors which are along $\mathbf{e}_s$ ($a^\phi = 0$) and tensors with vanishing off-diagonal components ($A^{s\phi} = A^{\phi s} = 0$). The non-zero components of covariant derivatives then read:

$$\nabla_s a^s = \partial_s a^s,\ \nabla_\phi a^\phi = \frac{\cos\psi}{x} a^s, \tag{9}$$

$$\nabla_s A^{ss} = \partial_s A^{ss},\ \nabla_s A^{\phi\phi} = \partial_s A^{\phi\phi} + 2\frac{\cos\psi}{x} A^{\phi\phi},\ \nabla_\phi A^{s\phi} = \nabla_\phi A^{\phi s} = -x\cos\psi A^{\phi\phi} + \frac{\cos\psi}{x} A^{ss}. \tag{10}$$

#### 1.2 Velocity field

The dynamics of the surface is described by the velocity field $\boldsymbol{v} = v^i \boldsymbol{e}_i + v_n \boldsymbol{n}$, with $v^i$ and $v_n$ the tangential and normal velocities. Due to axisymmetry, we assume that $v^\phi$ is zero. The strain-rate tensor is given by:

$$v_{ij} = \frac{\nabla_i v_j + \nabla_j v_i}{2} + C_{ij} v_n. \tag{11}$$

In the axisymetric shell geometry the only non zero components of the strain-rate tensor read, using Eq. 9:

$$v^s_s = \partial_s v^s + v_n C^s_s, \tag{12}$$

$$v^\phi_\phi = \frac{\cos\psi}{x} v^s + C^\phi_\phi v_n. \tag{13}$$

#### 1.3 Nematic field

We introduce a surface nematic field $Q_{ij}$ which accounts for anisotropic orientation of actomyosin fibers. The tensor field $Q_{ij}$ is symmetric and traceless. Due to the surface axisymmetry, off-diagonal components $Q_{s\phi} = Q_{\phi s}$ vanish, which implies that the nematic field is parametrized by the single quantity $q = Q^\phi_\phi = -Q^s_s$.

For simplicity we consider that the nematic field minimizes the following effective Frank-Landau-de Gennes free energy:

$$F_Q = \int_S \mathrm{d}S \left( \frac{k}{2} (\nabla_i Q^{jk})(\nabla^i Q_{jk}) - \frac{a}{4} Q_{ij} Q^{ij} + \frac{a}{16 S^2} (Q_{ij} Q^{ij})^2 \right), \tag{14}$$

where $dS$ is the differential surface area element of the surface $\mathcal{S}$, and $k$, $a$ and $S$ strictly positive constants. Introducing the normalized nematic tensor $\bar{Q}_{ij}$ defined by

$$Q_{ij} = S \bar{Q}_{ij}\ , \tag{15}$$

the effective free energy reads:

$$F_Q = S^2 \int_S \mathrm{d}S \left( \frac{k}{2} (\nabla_i \bar{Q}^{jk})(\nabla^i \bar{Q}_{jk}) - \frac{a}{4} \bar{Q}_{ij} \bar{Q}^{ij} + \frac{a}{16} (\bar{Q}_{ij} \bar{Q}^{ij})^2 \right), \tag{16}$$

The Euler-Lagrange equation on $q$ resulting from this free energy reads [2]:

$$\partial_s^2 \bar{q} = \frac{1}{2 l_c^2} \bar{q} (\bar{q}^2 - 1) + \frac{\cos\psi}{x} \left( 4 \frac{\cos\psi}{x} \bar{q} - \partial_s \bar{q} \right), \tag{17}$$

with $l_c = \sqrt{k/a}$ the correlation length of the nematic field, $\bar{q} = \bar{Q}^\phi_\phi = -\bar{Q}^s_s$ and:

$$q = S\bar{q}. \tag{18}$$

Using Eq. 18, the normalized profile of $\bar{q}$ can be solved numerically and the ordering strength $S$ adjusted to obtain the order profile $q(s)$. The length $l_c$ determines the size of topological defect cores at the aboral poles for all shapes, and at the oral pole for closed shapes. At distances large compared to $l_c$ away from the poles, the order relaxes to $q = \pm S$. Here we consider solutions relaxing to $q = S$, corresponding to circumferential order.

##### 1.4 Force and torque balance

As the ectoderm in cnidarians has a non-negligible thickness with respect to its transverse extension, we expect that its mechanics can not be simply describe by a purely two-dimensional stress tensor. pMLC stainings indicate that myosin has different distribution and spatial patterns on the basal side of the ectoderm/endoderm and apical side of the ectoderm (Fig. 2); we therefore expect that actomyosin contractile stresses exerted on these two surfaces differ. This differential stress profile along the tissue apico-basal axis generates bending moments on the mid-surface [2]. We therefore introduce the tension tensor $t_{ij}$, that accounts for the sum of stresses acting along the apico-basal axis, and the bending moment tensor $m_{ij}$, which accounts for differential stresses across the tissue apico-basal direction. For simplicity, we assume that there is no pressure difference between the regions inside and the outside of the tissue, even for species without oral opening, and more generally that no external forces or torques act on the surface, except at its boundary. The force balance reads, when projected on the tangent and the normal to the surface [3]:

$$\nabla_k t^{ki} + C^i_k t^k_n = 0, \tag{19}$$

$$\nabla_k t^k_n - C_{kl} t^{kl} = 0, \tag{20}$$

where $t^{ij}$ and $t^i_n$ are respectively the tangential and normal components of the tension tensor. Inertial terms are neglected as we expect tissue dynamics to occur at low Reynolds numbers.

The balance of torques acting on the surface reads:

$$t^i_n = \nabla_k \bar{m}^{ki}, \tag{21}$$

$$\epsilon_{kl} t^{kl} = C_{kl} m^{kl}, \tag{22}$$

with $\bar{m}^{ij} = -m^{ik} \epsilon_k{}^j$ the rotated bending moment tensor.

In the axisymmetric coordinates considered here, we take $t^{ij}$ and $\bar{m}^{ij}$ to have vanishing off-diagonal components, $t^{\phi}_n$ vanishes and the force and torque balance equations become: [2]:

$$\partial_s t^s_s = \frac{\cos\psi}{x}(t^{\phi}_{\phi} - t^s_s) - C^s_s t^s_n\,, \tag{23}$$

$$\partial_s t^s_n = C^{\phi}_{\phi}(t^{\phi}_{\phi} - t^s_s) + C_k{}^k t^s_s - \frac{\cos\psi}{x} t^s_n\,, \tag{24}$$

$$\partial_s \bar{m}^s_s = \frac{\cos\psi}{x}(\bar{m}^{\phi}_{\phi} - \bar{m}^s_s) + t^s_n\,. \tag{25}$$

##### 1.5 Constitutive equations

We describe the surface of cnidarian larvae as a viscoelastic anisotropic active two-dimensional fluid, where tensions and bending moments divide into three contributions:

$$t_{ij} = t^e_{ij} + t^v_{ij} + t^a_{ij}, \tag{26}$$

$$\bar{m}_{ij} = \bar{m}^e_{ij} + \bar{m}^v_{ij} + \bar{m}^a_{ij}, \tag{27}$$

where $t^e_{ij}$, $\bar{m}^e_{ij}$ correspond to elastic tensions and bending moments, $t^v_{ij}$ and $\bar{m}^v_{ij}$ to viscous dissipative tensions and bending moments, and $t^a_{ij}$ and $\bar{m}^a_{ij}$ are the active tension and bending moments. We assume that the elastic contributions follow the constitutive relations:

$$t^e_{ij} = 2Ku g_{ij} - 2\kappa C^k_k \left( C_{ij} - \frac{C^k_k}{2} g_{ij} \right), \tag{28}$$

$$\bar{m}^e_{ij} = 2\kappa C^k_k g_{ij}. \tag{29}$$

The first term in the tension tensor is an area elasticity term, with $K$ the area elastic modulus and $u = (\sqrt{g} - \sqrt{g_0})/\sqrt{g_0}$ the area strain, where $g = \det g_{ij}$ is the determinant of the metric tensor and $g_0$ the determinant of the metric tensor of the reference surface. The Lagrangian derivative of the area strain (i.e. its time derivative along points moving with the flow on the surface, denoted $Du/Dt$) then follows the dynamics [2]:

$$\frac{Du}{Dt} = (1+u)v_k^k. \tag{30}$$

The tension and bending moment proportional to $\kappa$, the bending modulus, account for bending elasticity of the tissue. Their expressions can be obtained from the Helfrich free energy [3]. In the following we consider a spatially varying profile of bending rigidity:

$$\kappa(s) = \kappa_0 + \delta\kappa(s) \tag{31}$$

with $\kappa_0$ a reference, uniform bending rigidity.

We consider shear, bulk and bending viscous effects leading to the following contributions:

$$t_{ij}^v = 2\eta\left(v_{ij} - \frac{v_k^k}{2}g_{ij}\right) + \eta_b v_k^k g_{ij}\ , \tag{32}$$

$$\bar{m}_{ij}^v = \eta_{cb}\frac{\mathrm{D}C_k^k}{\mathrm{Dt}}g_{ij}, \tag{33}$$

with $\eta$, $\eta_b$ and $\eta_{cb}$ the shear, bulk and bending viscosity and the corotational time derivative of the curvature tensor

$$\frac{\mathrm{D}C^{ij}}{\mathrm{Dt}} = -\nabla^i(\partial^j v_n) - v_n C_k^i C^{kj} + v_k\nabla^k C^{ij} + \omega_n\left(\epsilon^{ik}C_k^j + \epsilon^{jk}C_k^i\right), \tag{34}$$

with $\omega_n = \frac{1}{2}\epsilon^{ij}\nabla_i v_j$. The term $\frac{\mathrm{D}C_k^k}{\mathrm{Dt}}$ in Eq. 33 then reads in the coordinates used here:

$$\frac{\mathrm{D}C_k^k}{\mathrm{Dt}} = -\partial_s^2 v_n - \frac{\cos\psi}{x}\partial_s v_n - \left((C_s^s)^2 + (C_\phi^\phi)^2\right)v_n + v_s\partial_s C_k^k. \tag{35}$$

The active tension and bending moment tensors are given by:

$$t_{ij}^a = \zeta g_{ij} + \zeta_n Q_{ij}\ , \tag{36}$$

$$\bar{m}_{ij}^a = \zeta_c g_{ij} + \zeta_{cn} Q_{ij}, \tag{37}$$

with $\zeta$ the active isotropic tension, $\zeta_n$ the active nematic tension, $\zeta_c$ the active isotropic bending moment and $\zeta_{cn}$ the active nematic bending moment. In the following, we set $\zeta = 0$. Indeed, we note that the active isotropic tension results in a shift in the area deformation field, $u \to u + \zeta/(2K)$, in the isotropic elastic part of the tension tensor Eq. 28. At steady-state, let's assume that a solution of the force balance equations for a given profile $\zeta(s)$ can be obtained for the surface shape $\mathbf{X}(s)$ and area strain field $u(s) = u_s(s)$. In that case, a solution of the force balance equations with the same shape $\mathbf{X}(s)$, area strain field $u_s'(s) = u_s(s) + \zeta(s)/(2K)$ also exists, taking $\zeta = 0$ in the active isotropic tension, Eq. 36. In the limit of large $K$ that we take here, $u_s'(s) \simeq u_s(s)$ and the isotropic tension $\zeta$ has no effect.

#### 1.6 Tension and bending moment at the open boundary

For species with an open oral geometry, boundary conditions have to be specified on open side of the surface. We set the following conditions for the boundary tensions and bending moments:

$$t_s^s(L) = \left[2\pi k_P\frac{x(L)-r}{x(L)} + \frac{k_\psi}{2}\frac{(\psi(L)-\psi_r)^2}{x(L)}\right]\cos\psi(L), \tag{38}$$

$$t_n^s(L) = \left[2\pi k_P\frac{x(L)-r}{x(L)} + \frac{k_\psi}{2}\frac{(\psi(L)-\psi_r)^2}{x(L)}\right]\sin\psi(L), \tag{39}$$

$$\bar{m}_s^s(L) = -k_\psi(\psi(L)-\psi_r)\ , \tag{40}$$

where we take large values of constants $k_P$ and $k_\psi$ in order to constrain the value of the opening radius $x(L)$ to $r$, and of the opening angle $\psi(L)$ to $\psi_r$.

#### 2 Numerical simulations of cnidarian larval shapes

In this section, we describe numerical simulations of the active surface model described in section 1.

##### 2.1 Initial configuration

Simulations are initiated by considering a spherical cap:

$$\hat{x}(s_0) = \sin s_0, \tag{41}$$
$$\hat{z}(s_0) = 1 - \cos s_0, \tag{42}$$

where $s_0$ varies in $[0, 0.7\pi]$ for open shapes and in $[0, \pi]$ for closed shapes. Here $\hat{x} = x/R_0$ and $\hat{z} = z/R_0$ are normalised spatial coordinates. This also defines the reference surface with metric $g_0$.

##### 2.2 Numerical solving

To obtain stationary shapes for different oral geometries and active profiles, we simulated the dynamics of the open or closed active shell model with initial conditions defined in section 2.1. Numerical solving is performed by adapting the algorithm that has been described in Ref. [2]. We present here the main steps of the shape dynamics algorithm and refer to Ref. [2] for more details. The code used in this study is available at the repository https://github.com/DianaKhoromskaia/EpithelialShell/tree/open_boundary for open shapes and https://github.com/DianaKhoromskaia/EpithelialShell/tree/multiple_profiles for closed shapes.

**Determination of the nematic field.** To compute the velocity field $(v^s, v_n)$ at the surface of the shell at a time $t$, the nematic field is first determined by solving Eq. 17 with boundary conditions $\bar{q}(0) = 0$ and $\partial_s \bar{q}(0) = \partial_s \bar{q}(L) = 0$, and for closed surfaces, $\bar{q}(L) = 0$.

**Numerical solution of the mechanical equilibrium.** Replacing $t_{ij}$, $t_n^i$ and $\bar{m}_{ij}$ by their constitutive equations 26 and 27 in the mechanical equilibrium equations 23-25 and using relations Eqs. 6, 7 and 35, we obtain the following system of second order ODEs:

$$\begin{aligned}
\partial_s^2 v^s =& -\frac{\cos\psi}{x}\left(\partial_s v^s - \frac{\cos\psi}{x} v^s\right) - v_n \partial_s C_k^k - \frac{\eta - \eta_b}{\eta + \eta_b} C_s^s C_\phi^\phi v^s - \frac{1}{\eta + \eta_b}\left(\eta_b C_k^k + \eta(C_s^s - C_\phi^\phi)\right)\partial_s v_n \\
&+ \frac{1}{\eta + \eta_b}\left(-2K\partial_s u + (\partial_s \zeta_n) q + \zeta_n(\partial_s q) + 2\zeta_n q \frac{\cos\psi}{x} - C_s^s t_n^s - f_c \sin\psi\right) \\
&+ \frac{1}{\eta + \eta_b}\left((\partial_s \kappa) C_k^k (C_s^s - C_\phi^\phi) + 2\kappa C_s^s \partial_s C_k^k\right),
\end{aligned} \tag{43}$$

$$\partial_s t_n^s = C_\phi^\phi \left(t_\phi^\phi - t_s^s\right) + C_k^k t_s^s - \frac{\cos\psi}{x} t_n^s + f_c \cos\psi, \tag{44}$$

$$\partial_s \bar{m}_s^s = 2\zeta_{cn} q \frac{\cos\psi}{x} + t_n^s, \tag{45}$$

$$\partial_s^2 v_n = -\frac{\cos\psi}{x}\partial_s v_n - \left((C_\phi^\phi)^2 + (C_s^s)^2\right) v_n + v^s \partial_s C_k^k - \frac{1}{\eta_{cb}}\left(\bar{m}_s^s - 2\kappa C_k^k - \zeta_c + \zeta_{cn} q\right), \tag{46}$$

with

$$\begin{aligned}
t_s^s =& 2Ku - \kappa C_k^k \left(C_s^s - C_\phi^\phi\right) - \zeta_n q + (\eta + \eta_b)\partial_s v^s + (\eta_b - \eta)\frac{\cos\psi}{x} v^s \\
&+ \left(\eta_b C_k^k + \eta(C_s^s - C_\phi^\phi)\right) v_n ,
\end{aligned} \tag{47}$$

$$t_\phi^\phi - t_s^s = 2\left(\kappa C_k^k (C_s^s - C_\phi^\phi) + \zeta_n q + \eta\left(\frac{\cos\psi}{x} v^s - \partial_s v^s + (C_\phi^\phi - C_s^s) v_n\right)\right), \tag{48}$$

$$\bar{m}_\phi^\phi - \bar{m}_s^s = 2\zeta_{cn} q \,. \tag{49}$$

Here, an additional free parameter $f_c$ corresponding to an external force density $f_c \mathbf{e}_z$ has been introduced in the force balance equations Eqs. 43 and 44. This additional free parameter is used to numerically enforce the condition that the centroid of the shape does not move [2]. These equations are complemented with the boundary conditions $v^s(0) = t_n^s(0) = \partial_s v_n(0) = 0$ at the aboral pole, as a consequence of the fact that any continuous tangent vector field has to vanish at the pole of an axisymmetric surface. For closed surfaces, similar boundary conditions apply at the oral pole: $v^s(L) = t_n^s(L) = \partial_s v_n(L) = 0$. For surfaces which are open at the oral pole ($s = L$), boundary conditions are given by Eqs. 38-40.

We solve numerically the system of Eqs. 43-46. At the poles, $x = 0$ and to avoid numerical division by 0, one takes the following limits of Eqs. 43-46:

$$\text{At } s = 0: \ \partial_s^2 v^s = 0, \ \partial_s t_n^s = \frac{1}{2}(C_k^k t_s^s + f_c), \ \partial_s \bar{m}_s^s = 0, \ \partial_s^2 v_n = -\frac{1}{4}(C_k^k)^2 v_n - \frac{1}{2\eta_{cb}}(\bar{m}_s^s - 2\kappa C_k^k - \zeta_c) \ . \quad (50)$$

$$\text{At } s = L: \ \partial_s^2 v^s = 0, \ \partial_s t_n^s = \frac{1}{2}(C_k^k t_s^s - f_c), \ \partial_s \bar{m}_s^s = 0, \ \partial_s^2 v_n = -\frac{1}{4}(C_k^k)^2 v_n - \frac{1}{2\eta_{cb}}(\bar{m}_s^s - 2\kappa C_k^k - \zeta_c) \ . \quad (51)$$

where the limit in $s = L$ only applies to closed surfaces, as for open surfaces there is no singularities at $s = L$. After calculation of the velocity field $\boldsymbol{v} = v^s \boldsymbol{e}_s + v_n \boldsymbol{n}$, the surface position is updated using a Euler scheme:

$$\boldsymbol{X}(\phi, s, t + \mathrm{d}t) = \boldsymbol{X}(\phi, s, t) + \boldsymbol{v} dt \ . \quad (52)$$

Other geometrical fields are updated according to [2]:

$$\psi(s, t + dt) = \psi(s, t) + dt(-\partial_s v_n + v^s C_s^s) \ , \quad (53)$$

$$C_\phi^\phi(s, t + dt) = C_\phi^\phi(s, t) + dt\left(-\partial_s v_n \frac{\cos\psi}{x} - v_n (C_\phi^\phi)^2 + v^s \partial_s C_\phi^\phi\right) \ , \quad (54)$$

$$C_k^k(s, t + dt) = C_k^k(s, t) + dt \frac{1}{\eta_{cb}}\left(\bar{m}_s^s - 2\kappa C_k^k - \zeta_c + \zeta_{cn} q\right) \ , \quad (55)$$

$$\partial_s C_k^k(s, t + dt) = \partial_s C_k^k(s, t) + dt \frac{1}{\eta_{cb}}\left(t_n^s + 2\zeta_{cn} q \frac{\cos\psi}{x} + \partial_s(-2\kappa C_k^k - \zeta_c + \zeta_{cn} q\right) \ , \quad (56)$$

where Eqs. 53, 54 come from definitions of geometrical quantities, Eq. 55 from the constitutive equation for the bending moment (Eq. 27) and Eq. 56 from the torque balance equation 25.

The area strain field $u$ is updated according to:

$$u(s, t + \mathrm{d}t) = u(s, t) + (1 + u) v_k^k \mathrm{d}t, \quad (57)$$

which follows from Eq. 30. Following displacement of the shape, the new arclength coordinate is updated according to

$$s' = s + r_s dt \ , \quad (58)$$

$$\partial_s r_s = v_s^s = \partial_s v_s + v_n C_s^s \ . \quad (59)$$

We consider that the shape reaches steady-state once the maximum value of normalised velocity $v_n$ on the surface becomes less than $10^{-4}$.

#### 2.3 Computation of the apical shape

In order to compare simulation results to actual apical shapes of various species, the apical shape is reconstructed from the simulated mean shell shape $(x(s), z(s))$ and the animal ectoderm thickness profile $h(s)$. The basal and apical shapes are obtained as follows:

$$(x_{bas}(s), z_{bas}(s)) = (x(s), z(s)) - \frac{h(s)}{2}\boldsymbol{n}(s), \quad (60)$$

$$(x_{ap}(s), z_{ap}(s)) = (x(s), z(s)) + \frac{h(s)}{2}\boldsymbol{n}(s). \quad (61)$$

We show in Fig. 3F and Fig. S4C the effect on the apical shape of a thickness profile of the form:

$$\frac{h(s)}{R_0} = 0.2 + \frac{0.3}{1 + \exp(\frac{s-s_0+0.3}{50L})} - \frac{0.3}{1 + \exp(\frac{s-s_0-0.3}{50L})}\ , \tag{62}$$

for different values of $s_0$ on an spherical cap of opening angle $0.8\pi$ (Fig. 3F) or a sphere (Fig. S4C). To compare simulated and experimental shapes we used experimental thickness profiles.

### 3 Parameters and model fitting

Parameters value for simulations of different species are summarized in Tables 1-3. To obtain adimensional equations, we set $R_0$ as unit of length, $\kappa_0/R_0^2$ as unit of tension and $\eta R_0^3/\kappa_0$ as unit of time. This leaves eight free dimensionless parameters remaining in our system: $l_c/R_0$, $KR_0^2/\kappa_0$, $\eta_b/\eta$, $\eta_{cb}/\eta$, $k_P R_0^2/\kappa_0$, $k_\psi R_0/\kappa_0$, $r/R_0$ and $\psi_r$.

#### 3.1 Passive surface parameters

We set $KR_0^2/\kappa_0 = k_P R_0^2/\kappa_0 = k_\psi R_0/\kappa_0 = 200$. These numerical values are set large enough to effectively constrain the area of the shape and its oral geometry, such that their exact value is not essential. As we consider steady-state solutions, the values of viscosities $\eta_b/\eta$, $\eta_{cb}/\eta$ are chosen for convenience and do not influence our results. The value of $l_c/R_0 = 0.1$ is chosen so that the size of the defect at the aboral pole is small enough, such that the nematic order profile $\bar{q}$ relaxes quickly to its homogeneous value at $s \simeq 0.25$. Indeed, for $s > 0.25$ experimental profiles of basal order are roughly homogeneous (Fig. 2C).

The value of $r/R_0$ and $\psi_r$ are fixed by the experimental oral geometry of different species, at different stages of development (Table 1). We use experimental values obtained from average shapes in simulations (Table S6). In some cases, we round up experimental values to obtain simulation parameters, for simplicity.

| | Nematostella | Ctrl JNK-in-8 1hpf | JNK-in-8 1hpf | Aiptasia | Galaxea | Hydractinia | Cladonema |
|---|---|---|---|---|---|---|---|
| $l_c/R_0$ | 0.1 | 0.1 | 0.1 | 0.1 | 0.1 | 0.1 | 0.1 |
| $KR_0^2/\kappa_0$ | 200 | 200 | 200 | 200 | 200 | 200 | 200 |
| $\eta_b/\eta$ | 0.5 | 0.5 | 0.5 | 0.5 | 0.5 | 0.5 | 0.5 |
| $\eta_{cb}/\eta$ | 1 | 1 | 1 | 1 | 0.1 | 0.1 | 0.1 |
| $k_P R_0^2/\kappa_0$ | 200 | 200 | 200 | 200 | 200 | No mouth | No mouth |
| $k_\psi R_0/\kappa_0$ | 200 | 200 | 200 | 200 | 200 | No mouth | No mouth |
| $r/R_0$ | G: 0.32<br>EP: 0.27<br>P: 0.29<br>Myo.: 0.43 | G: 0.31<br>EP: 0.25<br>P: 0.27 | G: 0.31<br>EP: 0.29<br>P: 0.34 | G: 0.83<br>EP: 0.67<br>P: 0.56 | 0.22 | No mouth | No mouth |
| $\psi_r$ | $\pi$ | $\pi$ | G: $\pi$<br>EP: $0.9\pi$<br>P: $0.8\pi$ | G: $0.6\pi$<br>EP: $0.7\pi$<br>P: $0.6\pi$ | $0.8\pi$ | No mouth | No mouth |

Table 1: Simulation parameters for different species (parameters for a passive surface). Values for $r/R_0$ and $\psi_r$ are determined according to experimental measurements in Table S6. G accounts for *Gastrula*, EP for *Early Planula*, P for *Planula* and Myo. for *MyosinDN*.

#### 3.2 Parameters for model exploration

We give below parameter choices to study the individual effect of mechanical modules for open (Fig. 3B-E, 5B) and closed (Fig. S4C) surfaces.

- Active basal nematic tension (Fig. 3B, S4C, Movie S1): the maximal value of the order $q = Q^{\phi}_{\phi} = S\bar{Q}^{\phi}_{\phi}$ is varied by varying $S$, with constant active nematic tension $\zeta_n R_0^2/\kappa_0 = 8$ and bending moment $\zeta_{cn} R_0/\kappa_0 = 4/3$, with no isotropic tension or bending moment ($\zeta R_0^2/\kappa_0 = \zeta_c R_0/\kappa_0 = 0$), and with a uniform bending modulus ($\delta\kappa(s)/\kappa_0 = 0$). The value of the nematic coherence length is set to $l_c/R_0 = 0.1$.

- Oral geometry (Fig. 3C, 5B, Movie S2): simulations are performed for passive open surfaces ($\zeta = \zeta_c = \zeta_n = \zeta_{cn} = 0$) with uniform bending modulus ($\delta\kappa(s)/\kappa_0 = 0$) for different oral radii and angles. In Fig. 3C, $\psi_r$ varies keeping $r/R_0 = \sin 0.8\pi$, while in Fig. 5B both $r/R_0$ and $\psi_r$ vary.

- Aboral gradient of isotropic bending moment (Fig. 3D, S4C, Movie S3): simulations are performed with a step function of the uniform active bending moment $\zeta_c R_0/\kappa_0 \neq 0$ for $s < 0.2L$ and $\zeta_c R_0/\kappa_0 = 0$ for $s > 0.2L$. The value of $\zeta_c R_0/\kappa_0$ for $s < 0.2L$ is reported on the horizontal axis. Other active tensions or bending moments vanishing (($\zeta = \zeta_n = \zeta_{cn} = 0$), and with a uniform bending modulus ($\delta\kappa(s)/\kappa_0 = 0$).

- Increased aboral bending rigidity (Fig. 3E, S4C, Movie S4): simulations are performed with a non-uniform profile of bending modulus $\delta\kappa(s)/\kappa_0 = \frac{1}{1+\exp\left(\frac{s-l_\kappa}{100L}\right)}$ for different values of $l_\kappa$ and for passive surfaces ($\zeta = \zeta_c = \zeta_n = \zeta_{cn} = 0$).

In these simulations, we set $KR_0^2/\kappa_0 = 200$, $\eta_b/\eta = 1$, $\eta_{cb}/\eta = 1$. For open shapes we also set $k_P R_0^2/\kappa_0 = k_\psi R_0/\kappa_0 = 200$. For open surfaces (Fig. 3B-E) we chose by default $r/R_0 = \sin 0.8\pi$ and $\psi_r = 0.8\pi$.

#### 3.3 Determination of active profiles from experimental measurements

In this section we describe our rationale to obtain the profiles of active tensions magnitude $\zeta_n$, $\zeta_c$, $\zeta_{cn}$ (Eqs. 36-37) for various species simulated in Figs. 4, 5, S7, S9, S10.

##### 3.3.1 Tension and bending moment of active bilayer

Here we discuss active tensions and bending moments arising from apical and basal surface subjected to the active tension $t^{ap}_{ij}$, $t^{bas}_{ij}$. For small curvature $hC^k_k \ll 1$, and for apical and basal surfaces under active tensions $t^{ap}_{ij}$ and $t^{bas}_{ij}$, the resulting tension and bending moment tensor acting on the mean surface are:

$$t^a_{ij} = t^{ap}_{ij} + t^{bas}_{ij} , \tag{63}$$

$$\bar{m}^a_{ij} = \frac{h}{2}\left(t^{ap}_{ij} - t^{bas}_{ij}\right) . \tag{64}$$

with $h$ the distance between the apical and basal surfaces, corresponding to the ectoderm thickness. We consider that the active tension tensor of the apical and basal surface are given by:

$$t^{ap}_{ij} = \zeta^{ap} g_{ij} + \zeta^{ap}_n Q^{ap}_{ij} , \tag{65}$$

$$t^{bas}_{ij} = \zeta^{bas} g_{ij} + \zeta^{bas}_n Q^{bas}_{ij} . \tag{66}$$

where the metric tensor of the apical and basal surface is approximated by $g_{ij}$, for small curvature, and $Q^{ap}_{ij}$, $Q^{bas}_{ij}$ are the nematic tensors of the apical and basal surfaces. $\zeta^{ap}$, $\zeta^{bas}$ are the magnitude of the isotropic tensions on the apical and basal surfaces, and $\zeta^{ap}_n$, $\zeta^{bas}_n$ the magnitude of nematic tension on the apical and basal surfaces. We expect these magnitude to depend on the concentration of myosin on each surface. Therefore, we postulate that:

$$\zeta^{ap} = \beta^{ap} I^{ap} , \tag{67}$$

$$\zeta_n^{ap} = \beta_n^{ap} I^{ap} , \tag{68}$$
$$\zeta^{bas} = \beta^{bas} I^{bas} , \tag{69}$$
$$\zeta_n^{bas} = \beta_n^{bas} I^{bas} , \tag{70}$$

with $I^{ap}$, $I^{bas}$ the normalised pMLC fluorescence intensities measured on the apical and basal surfaces (Fig. 2B), and $\beta^{ap}$, $\beta_n^{ap}$, $\beta^{bas}$ and $\beta_n^{bas}$ proportionality coefficients. Overall, we arrive at the following expressions:

$$t_{ij}^a = (\zeta^{ap} + \zeta^{bas}) g_{ij} + \zeta_n^{ap} Q_{ij}^{ap} + \zeta_n^{bas} Q_{ij}^{bas} , \tag{71}$$
$$\bar{m}_{ij}^a = \frac{h}{2} (\zeta^{ap} - \zeta^{bas}) g_{ij} + \frac{h}{2} \zeta_n^{ap} Q_{ij}^{ap} - \frac{h}{2} \zeta_n^{bas} Q_{ij}^{bas} . \tag{72}$$

##### 3.3.2 Relating tension and bending moment profiles to experimental measurements

We now introduce the following simplifications:

- As most species (with the exception of *Hydractinia*) have small apical nematic order (Fig. 2C), we take in the following $Q_{ij}^{ap} = 0$.
- As discussed in section 1.5, the active isotropic total tension has no effect in the case of a nearly incompressible surface as we consider here, and we therefore do not consider further this contribution.
- Since the basal pMLC intensity profiles are relatively homogeneous (Fig. 2B), we take them to be uniform. Uniform, isotropic contributions to $\bar{m}_{ij}$ do not change the active surface flows and deformation, since such contributions disappear from the torque balance equation 25 and do not contribute to the boundary condition 40, if $k_\psi$ is large enough. We therefore do not consider further the contribution in $\zeta^{bas} g_{ij}$ in Eq. 72.

With these simplifications, we obtain:

$$t_{ij}^a = \zeta_n^{bas} Q_{ij}^{bas} , \tag{73}$$
$$\bar{m}_{ij}^a = \frac{h}{2} \zeta^{ap} g_{ij} - \frac{h}{2} \zeta_n^{bas} Q_{ij}^{bas} . \tag{74}$$

We identify the nematic field $Q_{ij}$ introduced in section 1.3 with the basal nematic field $Q_{ij}^{bas}$. Using Eq. 15 to introduce the normalised nematic tensor $\bar{Q}_{ij}$, the normalized active tension and bending moment tensors $\hat{t}_{ij}^a = t_{ij}^a R_0^2 / \kappa_0$, $\hat{\bar{m}}_{ij}^a = \bar{m}_{ij}^a R_0 / \kappa_0$ are given by:

$$\hat{t}_{ij}^a = \hat{\zeta}_n \bar{Q}_{ij} , \tag{75}$$
$$\hat{\bar{m}}_{ij}^a = \hat{\zeta}_c g_{ij} + \hat{\zeta}_{cn} \bar{Q}_{ij} . \tag{76}$$

with the normalized active tension magnitudes $\hat{\zeta}_n$, $\hat{\zeta}_c$, $\hat{\zeta}_{cn}$, given by:

$$\hat{\zeta}_n(s) = \beta_n Z_n, \tag{77}$$
$$\hat{\zeta}_c(s) = \beta Z_c(s), \tag{78}$$
$$\hat{\zeta}_{cn}(s) = -\beta_n Z_{cn}, \tag{79}$$

with $\beta_n = \beta_n^{bas} R_0^2/\kappa_0$, $\beta = \beta^{ap} R_0^2 / \kappa_0$, and we have introduced the dimensionless profile functions $Z_n(s)$, $Z_c(s)$, $Z_{cn}(s)$. The functions $Z_n(s)$, $Z_c(s)$, $Z_{cn}(s)$ are evaluated based on experimental measurements:

$$Z_n = \left\langle I^{bas}(s) \tilde{Q}_\phi^{\phi, bas, exp}(s) \right\rangle_s , \tag{80}$$
$$Z_c(s) = \frac{h(s) I^{ap}(s)}{2R_0}, \tag{81}$$
$$Z_{cn} = \frac{\left\langle h(s) I^{bas}(s) \tilde{Q}_\phi^{\phi, bas, exp}(s) \right\rangle_s}{2R_0} . \tag{82}$$

Here, $h(s)$ is the profile of tissue thickness (Fig. S2C). $\tilde{Q}_{\phi}^{\phi,bas,exp}(s) = Q_{\phi}^{\phi,bas,exp}(s)A^{bas}(s)$ is the product of the experimentally measured average profile of basal nematic order (Fig. 2C), and $A^{bas}(s)$ the average scalar alignment parameter (Fig. S2D, Material and Methods). The nematic $\tilde{Q}_{\phi}^{\phi,bas,exp}(s)$ gives an overall measure of circumferential alignment obtained from the pMLC fluorescence intensity, used to evaluate the amplitude of order $S$. The averaging sign $\langle\cdot\rangle_s$ corresponds to average of experimental profiles over $s$, with $s$ values which are in general away from the aboral poles for all species, and away from the aboral and oral poles for species with closed shapes. $Z_c$ profiles were obtained by fitting experimental data with a sigmoid function for *Nematostella*, *Aiptasia* and *Cladonema*:

$$Z_c(s) = \frac{Z_0}{1 + \exp\left(\frac{s-l_a}{wL}\right)} + \bar{Z}_c, \tag{83}$$

and by an inverse exponential function for *Galaxea* and *Hydractinia*:

$$Z_c(s) = Z_0 \exp\left(-\frac{s}{wL}\right) + \bar{Z}_c. \tag{84}$$

The value of $Z_n$, $Z_{cn}$, $Z_0$, $l_a$, $w$ for each species is reported in Table 2. In practice we set $\bar{Z}_c = 0$ when performing simulations, as mechanical equilibrium is invariant by the addition of a uniform isotropic bending moment.

Eqs. 77-79 indicate that the magnitude of isotropic active bending moments depends on $\beta$, and the magnitude of anisotropic active tension and bending moments depend on $\beta_n$. Since, as discussed above, uniform and isotropic contributions to $\bar{m}_{ij}$ do not affect the surface shape, only spatial variations of $\hat{\zeta}_c(s)$ are relevant; these spatial variations depend on the apical myosin profiles, which consist mostly of an increase towards the aboral side (Fig. 2). Therefore, we associate $\beta$ to the “aboral isotropic active bending moment" module, and $\beta_n$ to the “active basal nematic tension" module.

##### 3.4 Bending modulus profile

We assume that as a result of the gradient of ectoderm thickness, the tissue has a spatial profile of bending modulus $\kappa(s)$. We choose the following profile of dimensionless bending modulus:

$$\hat{\kappa}(s) = 1 + \frac{\hat{\kappa}_{ab} - 1}{1 + \exp\left(\frac{s-l_\kappa}{w_\kappa L}\right)}, \tag{85}$$

with $w_\kappa = 100$. With this profile, the normalized bending modulus sharply transitions from $\hat{\kappa}_{ab}$ when $s < l_\kappa$ to 1 for $s > l_\kappa$. The parameter $l_\kappa$ is determined from the average ectoderm thickness profile for each species as the value of $s$ such that

$$h(l_\kappa) - h_{min} = 0.9\left(h_{max} - h_{min}\right), \tag{86}$$

with $h_{min}$ and $h_{max}$ respectively the lowest and highest value of the ectoderm thickness. The value of $l_\kappa$ for each species is reported in Table 2.

##### 3.5 Determination of species-dependent parameters

To determine the value of the free parameters $\beta$, $\beta_n$ and $\hat{\kappa}_{ab}$, simulated steady-state shapes $(\hat{x}(s), \hat{z}(s))$ (with $s \in [0, L]$) are compared to experimentally measured shapes $(\hat{x}^*(s), \hat{z}^*(s))$ (with $s \in [0, L^*]$, with $L^*$ the total length of the experimental contour). We introduce the following error function:

$$\Delta\left[(\hat{x}, \hat{z}), (\hat{x}^*, \hat{z}^*)\right] = \int_0^L \min_{s' \in [0, L^*]} \left[(\hat{x}(s) - \hat{x}^*(s'))^2 + (\hat{z}(s) - \hat{z}^*(s'))^2\right] \mathrm{d}s, \tag{87}$$

where we choose the origin of $\hat{z}$ and $\hat{z}^*$ as the middle point between oral and aboral extremities and with $\min_{s' \in [0,L^*]} \left[(\hat{x}(s) - \hat{x}^*(s'))^2 + (\hat{z}(s) - \hat{z}^*(s'))^2\right]$ the minimal distance between the simulated shape at position $s$

| | Nematostella | Ctrl JNK-in-8 1hpf | JNK-in-8 1hpf | Aiptasia | Galaxea | Hydractinia | Cladonema |
|---|---|---|---|---|---|---|---|
| $l_a/L$ | G: 0.13<br>EP: 0.14<br>P: 0.1<br>Myo.: 0.11 | G: 0.2<br>EP: 0.25<br>P: 0.2 | G: 0.17<br>EP: 0.26<br>P: 0.31 | G: 0.15<br>EP: 0.29<br>P: 0.3 | not applicable | not applicable | 0.04 |
| $Z_0$ | G: 0.5<br>EP: 0.7<br>P: 1.3<br>Myo.: 1.1 | G: 0.18<br>EP: 0.24<br>P: 0.39 | G: 0.86<br>EP: 1.1<br>P: 0.94 | G: 1.8<br>EP: 3.1<br>P: 2.9 | 1.8 | 1.5 | 5.2 |
| $w$ | G: 0.06<br>EP: 0.07<br>P: 0.07<br>Myo.: 0.06 | G: 0.08<br>EP: 0.06<br>P: 0.07 | G: 0.17<br>EP: 0.15<br>P: 0.09 | 0.08 | 0.45 | 0.12 | 0.12 |
| $Z_n$ | G: 0.05<br>EP: 0.15<br>P: 0.24<br>Myo.: 0.02 | G: 0.03<br>EP: 0.07<br>P: 0.11 | G: 0.02<br>EP: 0.08<br>P: 0.08 | G: 0.12<br>EP: 0.24<br>P: 0.3 | 0.11 | 0.16 | 0.01 |
| $Z_{cn}$ | G: 0.008<br>EP: 0.02<br>P: 0.03<br>Myo.: 0.00002 | G: 0.005<br>EP: 0.01<br>P: 0.01 | G: 0.004<br>EP: 0.01<br>P: 0.01 | G: 0.04<br>EP: 0.05<br>P: 0.04 | 0.009 | 0.02 | 0.003 |
| $l_\kappa/L$ | G: 0.16<br>EP: 0.1<br>P: 0.08<br>Myo.: 0.07 | G: 0.14<br>EP: 0.14<br>P: 0.13 | G: 0.12<br>EP: 0.23<br>P: 0.19 | G: 0.1<br>EP: 0.16<br>P: 0.2 | 0.04 | 0.03 | 0.07 |

Table 2: Table of parameter values for the active and bending modulus spatial profiles. G accounts for *Gastrula*, EP for *Early Planula*, P for *Planula* and Myo. for *MyosinDN*.

| | Nematostella | Aiptasia | Galaxea | Hydractinia | Cladonema |
|---|---|---|---|---|---|
| $R_0$ | WT+JNK+CTRL: 120$\mu m$<br>SQH: 110$\mu$m | Gastrula: 40$\mu m$<br>Early Planula: 40$\mu m$<br>Planula: 45$\mu m$ | 220$\mu$m | 90$\mu$m | 20$\mu$m |

Table 3: Magnitude of reference radius $R_0$ for different species, used for comparison between theory and experiment. See Table S6 for parameter measurements.

and the experimental shape. The experimental normalised shape functions $\hat{z}^*(s)$ and $\hat{x}^*(s)$ are obtained by normalising the coordinates $x^*(s)$, $z^*(s)$ of the average shape by the reference radius $R_0$ (Table 3). For *Galaxea*, *Hydractinia* and *Cladonema*, pMLC profiles are measured at planula stage, such that the value of $\beta$, $\beta_n$ and $\hat{\kappa}_{ab}$ are obtained by minimizing $\Delta\left[(\hat{x},\hat{z}),(\hat{x}^*,\hat{z}^*)\right]$ at this stage. For *Nematostella* and *Aiptasia*, pMLC intensity profiles are measured at three different stages (gastrula, early planula and planula). Considering that the free parameters do not change during development, we look for a single value of the fitting parameters that minimizes the average of $\Delta\left[(\hat{x},\hat{z}),(\hat{x}^*,\hat{z}^*)\right]$ over the three different stages of development. The optimal triplet $(\beta,\beta_n,\hat{\kappa}_{ab})$ is found by simulating shapes obtained for a range discrete values of the three parameters, and identifying the triplet that minimizes the fit criterion. Parameter values sampled to perform the fit are reported in Table 4.

| | Discrete values of $\beta$ | Discrete values of $\beta_n$ | Discrete values of $\hat{\kappa}_{ab}$ |
|---|---|---|---|
| *Nematostella* | $I(0,2,0.5)$ | $0, I(10,20,2)$ | $I(1,2,0.25)$ |
| *Aiptasia* | $I(0,2,0.5)$ | $I(0,50,5)$ | $I(1,3,0.5)$ |
| *Galaxea* | $I(0,4,1)$ | $0, I(10,40,5)$ | $I(1,5,0.5)$ |
| *Hydractinia* | $I(0,6,1)$ | $0, I(50,100,5)$ | $I(1,7,0.5)$ |
| *Cladonema* | $I(0,1,0.25)$ | $0, I(75,275,25)$ | $I(1,2,0.25)$ |

Table 4: Discrete values of $\beta$, $\beta_n$ and $\hat{\kappa}_{ab}$ used to perform the fit for each species. The notation $I(a,b,c)$ accounts for the sequence of number going from $a$ to $b$ with step $c$.

We first performed a fit simultaneously for the three stages of *Nematostella* wild-type, and controls of JNK-in-8 treated *Nematostella*, with fixed parameters $\beta_n$, $\beta$, $\hat{\kappa}_{ab}$. We found a best fit for $\beta_n = 14$, $\beta = 0$, $\hat{\kappa}_{ab} = 1$, which we show in Fig. 4C. This results corresponds to neither aboral gradient of isotropic active bending moment, nor increased aboral bending rigidity, acting in *Nematostella* in normal conditions.

Assuming that these parameters did not change in JNK-in-8 treated *Nematostella* compared to wild-type and control *Nematostella*, we found that in simulations JNK-in-8 treated *Nematostella* had increased shape polarity compared to control, but the increase was not as pronounced as in experiments (Fig.S9 E,F; blue boxes).

To clarify this point, we first asked whether the JNK-in-8 perturbation could have affected the value of $\beta$, while keeping $\hat{\kappa}_{ab} = 1$; corresponding to an aboral gradient of isotropic active bending moment, but no increased aboral bending rigidity. We performed a fit on the three stages of JNK-in-8 treated *Nematostella*, forcing $\hat{\kappa}_{ab} = 1$ and $\beta_n$ to its value obtained from the fit on *Nematostella* wild-type and control of JNK-in-8. We obtained the best fit parameter $\beta = 1$, different from 0, but simulated shapes were still strongly underestimating the shape polarity (Fig. S9E,F, red boxes).

We then instead asked whether JNK-in-8 treatment of *Nematostella* could have modified the value of $\hat{\kappa}$ while keeping $\beta = 0$; corresponding to an increased aboral bending rigidity, but no aboral gradient of isotropic active bending moment. We therefore performed a fit adjusting the value of $\hat{\kappa}$, while keeping the values of $\beta$ and $\beta_n$ obtained from the fit on *Nematostella* wild-type and control. Doing so, we obtain the best fit parameter $\hat{\kappa} = 1.25$, and observe improvement in the simulated shapes, with a clear change of the sign of the JNK-in-8 treated *Nematostella* planula mean shell polarity compared to control (Fig. S9E,F, black boxes). The simulated shape polarity was however still lower than the experimental shape polarity, indicating that other factors are at play. Overall, we conclude that increased aboral bending rigidity, rather than an aboral gradient of isotropic active bending moment, plays a role in setting the shape of JNK-in-8 treated *Nematostella*.

To further test this conclusion, and since we assume that increased bending rigidity is correlated to tissue thickening on the aboral side, we asked if the range of the increased thickness aboral domain and the magnitude of the thickness increase was correlated to shape polarity change, across individuals of JNK-in-8 treated *Nematostella*. We defined an increase thickness domain range $l_\kappa$ for each JNK-in-8 treated individual, by fitting the ectoderm thickness profile by a sigmoid function in the region $s \in [0, s_{min}]$, with $s_{min}$ the position of the minimum of thickness, and using the criterion (86) to compute $l_\kappa$ from the sigmoidal fitted thickness profile. We observed that the mean shell polarity indeed varied significantly with $l_\kappa/L$ across individuals. Interestingly the variation was non-linear with a maximum polarity at intermediate values of $l_\kappa$, in agreement with simulation results (Fig. S9D, left panel). In addition we also observed a correlation between the mean shell polarity and the ratio of maximum to minimum tissue thickness (Fig.S9D, middle panel); consistent with the idea that increased aboral tissue thickness leads to higher aboral bending rigidity and therefore more positive shape polarities. This is also consistent with the fit result that $\hat{\kappa}_{ab}$ increases following JNK-in-8 treatment, as in JNK-in-8 perturbations the aboral domain is thicker than in wild-type (Fig. 5E). Finally, we also found that shape polarity did not correlate with a measure of the gradient of aboral isotropic bending moment, $\Delta(h \times I^{ap}) = \max(h \times I^{ap}) - \min(h \times I^{ap})$; across JNK-in-8 treated individuals (Fig. S9D, right panel). Overall these results are consistent with the idea that increased bending rigidity, but not the aboral gradient of isotropic active bending moment, contributes in setting larval shapes.

Turning to other species larval shapes, we first performed fits imposing $\beta = 0$ and $\hat{\kappa}_{ab} = 1$, corresponding to neither aboral gradient of isotropic active bending moment, nor increased aboral bending rigidity. We obtained reasonable results for *Galaxea* and *Cladonema*, but did not capture well the shape polarities of *Aiptasia* and *Hydractinia* shapes (Fig. S7A, fourth column. Fig. S7C). We then performed fits imposing $\hat{\kappa}_{ab} = 1$ and leaving $\beta$ as a free parameter, but we did not notice an improvement in simulated shapes of *Aiptasia* and *Hydractinia* (Fig.S7A, third column, Fig. S7B). Finally, imposing $\beta = 0$ and adjusting $\beta_n$ and $\hat{\kappa}_{ab}$ resulted in reasonable agreement between experimental shapes and optimal simulated shapes (Fig. 4J, 5I, Fig. S7A, second column). We conclude that the aboral gradient of active isotropic bending moment is not necessary to explain cnidarian larvae shapes, but a gradient of bending rigidity is. Therefore, we then fitted all species with the assumption $\beta = 0$. The optimal values of $\beta_n$ and $\hat{\kappa}_{ab}$ (imposing $\beta = 0$) and optimal values of $\beta_n$ and $\beta$ (imposing $\hat{\kappa}_{ab} = 1$) are reported in Table 5.

| | | Nematostella WT + control of JNK-in-8 | JNK ($\beta_n$ set by Nematostella WT +control of JNK-in-8) | Aiptasia | Galaxea | Hydractinia | Cladonema |
|---|---|---|---|---|---|---|---|
| Fit with $\beta = 0$ | $\beta_n$ | 14 | 14 | 10 | 35 | 55 | 150 |
| | $\hat{\kappa}_{ab}$ | 1 | 1.25 | 2 | 4.5 | 4.5 | 1.75 |
| Fit with $\hat{\kappa}_{ab} = 1$ | $\beta_n$ | 14 | 14 | 5 | 35 | 50 | 175 |
| | $\beta$ | 0 | 1 | 0 | 3 | 6 | 0.75 |
| Fit with $\beta = 0$ and $\hat{\kappa}_{ab} = 1$ | $\beta_n$ | 14 | 14 | 5 | 35 | 50 | 125 |

Table 5: Fit parameter values for five species, assuming $\beta = 0$ (top, no gradient of isotropic active bending moment) or $\hat{\kappa}_{ab} = 1$ (middle, no increased aboral bending rigidity), or both $\beta = 0$ and $\hat{\kappa}_{ab} = 1$ (bottom, no gradient of isotropic active bending moment and no increased aboral bending rigidity).

### 4 Quantification of the relative contributions of different effects to the shape

We describe here a method to quantify the relative contributions of mechanical modules (originating from basal active nematic tension, aboral active isotropic bending moment, opening geometry, increased aboral bending rigidity and thickness profile) to the cnidarian larval shape, its aspect ratio and polarity.

We denote $X$ a scalar quantity characterizing the shape (here $X$ can correspond to the shape elongation or the shape polarity). $X^*$ denotes the value of $X$ for a reference shape where none of the mechanical modules are introduced. Here, the reference shape is taken as a sphere. For simplicity, we take $X^*$ as the origin of $X$, such that in what is next we made the transformation $X \to X - X^*$.

The value of $X$ obtained by combining $l$ distinct modules is written $X_{i_1,...,i_l}$, where $i_1, ..., i_l$ are distinct integers labelling different modules. The value of $X_{i_1,...,i_l}$ can be obtained by running simulations including these $l$ modules (Fig. S10). $X_{i_1,...i_l}$ is invariant by permutation of the indices, as all $l$ modules are applied simultaneously, so that their order does not matter. Therefore, in the following indices are taken in ascending order. We follow an approach related to many-body expansion used e.g. in liquid physics [1] and the exclusion-inclusion principle [4]. The value of $X$ for any combination of two or three modules can be written as

$$X_{i,j} = \Delta X_i + \Delta X_j + \Delta X_{i,j}, \tag{88}$$

$$X_{i,j,k} = \Delta X_i + \Delta X_j + \Delta X_k + \Delta X_{i,j} + \Delta X_{i,k} + \Delta X_{j,k} + \Delta X_{i,j,k}, \tag{89}$$

where

$$\Delta X_i = X_i, \tag{90}$$

$$\Delta X_{i,j} = X_{i,j} - \Delta X_i - \Delta X_j, \tag{91}$$

$$\Delta X_{i,j,k} = X_{i,j,k} - \Delta X_{i,j} - \Delta X_{i,k} - \Delta X_{j,k} - \Delta X_i - \Delta X_j - \Delta X_k. \tag{92}$$

Here, $\Delta X_i$ corresponds to the contribution to $X$ of the effect $i$ in isolation. $\Delta X_{i,j}$ corresponds to the contribution of the effects $i, j$ which is not explained by the sum of the individual contributions of $i$, $j$. $\Delta X_{i,j,k}$ is the contribution of the effects $i, j, k$ that can not be explained by the sum of the individual contributions $i, j, k$, or the contributions of the pairs $(i, j)$, $(j, k)$, $(i, k)$.

We can generalize this decomposition to the combination $c = (i_1, ..., i_l)$ of $l$ modules:

$$\Delta X_c = X_c - \sum_{k=1}^{l-1} \sum_{c' \in C_k(c)} \Delta X_{c'}, \tag{93}$$

where $C_k(c)$ is the ensemble formed by all ordered combinations of $k$ distinct modules present in $c$. $\Delta X_c$ represents the contribution to $X$ of combination $c$ that can not be explained by the sum of lower order combinations.

For one, two and three modules one can verify that the definitions introduced above are equivalent to:

$$\Delta X_i = X_i, \tag{94}$$

$$\Delta X_{i,j} = X_{i,j} - X_i - X_j, \tag{95}$$

$$\Delta X_{i,j,k} = X_{i,j,k} - X_{i,j} - X_{i,k} - X_{j,k} + X_i + X_j + X_k, \tag{96}$$

and for an arbitrary number of modules:

$$\Delta X_c = \sum_{k=1}^{l} (-1)^{l-k} \sum_{c' \in C_k(c)} X_{c'}. \tag{97}$$

This last relationship can be shown by induction; indeed assuming that $\Delta X_{c'} = \sum_{k=1}^{m} (-1)^{m-k} \sum_{c'' \in C_k(c')} X_{c''}$, for all $c' \in C_m(c)$ and all $m < l$, one then obtains, starting from Eq. 93:

$$\begin{aligned}
\Delta X_c &= X_c - \sum_{m=1}^{l-1} \sum_{c'' \in C_m(c)} \sum_{k=1}^{m} (-1)^{m-k} \sum_{c' \in C_k(c'')} X_{c'} \\
&= X_c - \sum_{m=1}^{l-1} \sum_{k=1}^{m} (-1)^{m-k} \binom{l-k}{m-k} \sum_{c' \in C_k(c)} X_{c'} \\
&= X_c - \sum_{k=1}^{l-1} \sum_{m=k}^{l-1} (-1)^{m-k} \binom{l-k}{m-k} \sum_{c' \in C_k(c)} X_{c'} \\
&= X_c - \sum_{k=1}^{l-1} \left[ (1-1)^{l-k} - (-1)^{l-k} \right] \sum_{c' \in C_k(c)} X_{c'}
\end{aligned}$$

$$= \sum_{k=1}^{l} (-1)^{l-k} \sum_{c' \in C_k(c)} X_{c'}. \tag{98}$$

Eq. 93 can be rewritten as a decomposition equation of $X_c$ into a sum over contribution of $k$ simultaneously introduced modules:

$$X_c = \sum_{k=1}^{l} \sum_{c' \in C_k(c)} \Delta X_{c'}, \tag{99}$$

which for three modules labelled $1, 2, 3$ read:

$$X_{1,2,3} = \Delta X_1 + \Delta X_2 + \Delta X_3 + \Delta X_{1,2} + \Delta X_{1,3} + \Delta X_{2,3} + \Delta X_{1,2,3} \; . \tag{100}$$

In order to assign an overall contribution of each module, we rewrite Eq. 99 as follows:

$$X_c = \sum_{i \in c} \delta X_i, \tag{101}$$

with

$$\delta X_i = \sum_{k=1}^{l} \frac{1}{k} \sum_{c' \in C_k^i(c)} \Delta X_{c'}, \tag{102}$$

the contribution to $X_c$ of module $i$, with $C_k^i(c)$ the set of combinations of $k$ modules of $c$, that include module $i$. For three modules labelled $1, 2, 3$, this reads:

$$X_{1,2,3} = \delta X_1 + \delta X_2 + \delta X_3 \; , \tag{103}$$

$$\delta X_1 = \Delta X_1 + \frac{1}{2}(\Delta X_{1,2} + \Delta X_{1,3}) + \frac{1}{3}\Delta X_{1,2,3} \; , \tag{104}$$

$$\delta X_2 = \Delta X_2 + \frac{1}{2}(\Delta X_{1,2} + \Delta X_{2,3}) + \frac{1}{3}\Delta X_{1,2,3} \; , \tag{105}$$

$$\delta X_3 = \Delta X_3 + \frac{1}{2}(\Delta X_{1,3} + \Delta X_{2,3}) + \frac{1}{3}\Delta X_{1,2,3} \; . \tag{106}$$

Here the contribution of $\Delta X_{1,2}$ is split equally between $\delta X_1$ and $\delta X_2$, the contribution of $\Delta X_{1,2,3}$ is split equally between $\delta X_1$, $\delta X_2$, $\delta X_3$, etc. More generally, going from Eq. 99 to 101, $\Delta X_{c'}$ is contributing a gain $\Delta X_{c'}/\mathrm{card}(c')$ to all modules within $c'$.

We now seek to express $\delta X_i$ in terms of sums of $X_{i_1,\ldots,i_k}$. Using Eq. 97, Eq. 102 can be rewritten as:

$$\begin{aligned}
\delta X_i &= \sum_{m=1}^{l} \frac{1}{m} \sum_{c'' \in C_m^i(c)} \sum_{k=1}^{m} (-1)^{m-k} \left( \sum_{c' \in C_k^i(c'')} X_{c'} + \sum_{c' \in C_{k+1}^i(c'')} X_{c' \setminus \{i\}} \right) \\
&= \sum_{m=1}^{l} \frac{1}{m} \left[ \sum_{k=1}^{m} (-1)^{m-k} \binom{l-k}{m-k} \sum_{c' \in C_k^i(c)} X_{c'} \right] + \sum_{m=2}^{l} \frac{1}{m} \left[ \sum_{k=1}^{m-1} (-1)^{m-k} \binom{l-k-1}{m-k-1} \sum_{c' \in C_{k+1}^i(c)} X_{c' \setminus \{i\}} \right] \\
&= \sum_{k=1}^{l} \left[ \sum_{m=k}^{l} \frac{(-1)^{m-k}}{m} \binom{l-k}{m-k} \right] \sum_{c' \in C_k^i(c)} X_{c'} + \sum_{k=1}^{l} \left[ \sum_{m=k}^{l} \frac{(-1)^{m-k+1}}{m} \binom{l-k}{m-k} \right] \sum_{c' \in C_k^i(c)} X_{c' \setminus \{i\}} \\
&= \sum_{k=1}^{l} A_k^l \sum_{c' \in C_k^i(c)} (X_{c'} - X_{c' \setminus \{i\}}),
\end{aligned} \tag{107}$$

with $A_k^l = \sum_{m=0}^{l-k} \frac{(-1)^m}{m+k}\binom{l-k}{m}$ and where we have used that $C_m(c') = C_m^i(c') \cup C_m(c'\backslash\{i\})$, $C_m(c'\backslash\{i\}) = \{c"\backslash\{i\} | c" \in C_{m+1}^i(c')\}$, $C_{m+1}^i(c") = \emptyset$ ($m$ being the number of elements in $c"$) and $X_\emptyset = 0$. Noticing that $A_k^l + A_{k+1}^l = A_k^{l-1}$, one can show by induction that

$$A_k^l = \frac{(k-1)!(l-k)!}{l!} = \frac{1}{l\binom{l-1}{k-1}}. \tag{108}$$

Then $\delta X_i$ reads

$$\delta X_i = \frac{1}{l}\sum_{k=1}^{l} \langle \delta X \rangle_i^{(k)}, \tag{109}$$

with

$$\langle \delta X \rangle_i^{(k)} = \frac{1}{\binom{l-1}{k-1}} \times \sum_{c' \in C_k^i(c)} (X_{c'} - X_{c'\backslash\{i\}}), \tag{110}$$

the average gain of $X$ generated by the addition of the module $i$ to $k$ other effects. For the case of three modules $1, 2, 3$, this corresponds to:

$$\delta X_1 = \frac{1}{3}\left[X_1 + \frac{(X_{1,2} - X_2) + (X_{1,3} - X_3)}{2} + (X_{1,2,3} - X_{2,3})\right], \tag{111}$$

$$\delta X_2 = \frac{1}{3}\left[X_2 + \frac{(X_{1,2} - X_1) + (X_{2,3} - X_3)}{2} + (X_{1,2,3} - X_{1,3})\right], \tag{112}$$

$$\delta X_3 = \frac{1}{3}\left[X_3 + \frac{(X_{1,3} - X_1) + (X_{2,3} - X_2)}{2} + (X_{1,2,3} - X_{1,2})\right]. \tag{113}$$

For the case of four modules $1, 2, 3, 4$, one has $X_c = \delta X_1 + \delta X_2 + \delta X_3 + \delta X_4$ with:

$$\begin{aligned}\delta X_1 =& \frac{1}{4}\left[X_1 + \frac{(X_{1,2} - X_2) + (X_{1,3} - X_3) + (X_{1,4} - X_4)}{3}\right.\\ &\left.+\frac{(X_{1,2,3} - X_{2,3}) + (X_{1,2,4} - X_{2,4}) + (X_{1,3,4} - X_{3,4})}{3} + (X_{1,2,3,4} - X_{2,3,4})\right],\end{aligned} \tag{114}$$

$$\begin{aligned}\delta X_2 =& \frac{1}{4}\left[X_2 + \frac{(X_{1,2} - X_1) + (X_{2,3} - X_3) + (X_{2,4} - X_4)}{3}\right.\\ &\left.+\frac{(X_{1,2,3} - X_{1,3}) + (X_{1,2,4} - X_{1,4}) + (X_{2,3,4} - X_{3,4})}{3} + (X_{1,2,3,4} - X_{1,3,4})\right],\end{aligned} \tag{115}$$

$$\begin{aligned}\delta X_3 =& \frac{1}{4}\left[X_3 + \frac{(X_{1,3} - X_1) + (X_{2,3} - X_2) + (X_{3,4} - X_4)}{3}\right.\\ &\left.+\frac{(X_{1,2,3} - X_{1,2}) + (X_{1,3,4} - X_{1,4}) + (X_{2,3,4} - X_{2,4})}{3} + (X_{1,2,3,4} - X_{1,2,4})\right],\end{aligned} \tag{116}$$

$$\begin{aligned}\delta X_4 =& \frac{1}{4}\left[X_4 + \frac{(X_{1,4} - X_1) + (X_{2,4} - X_2) + (X_{3,4} - X_3)}{3}\right.\\ &\left.+\frac{(X_{1,2,4} - X_{1,2}) + (X_{1,3,4} - X_{1,3}) + (X_{2,3,4} - X_{2,3})}{3} + (X_{1,2,3,4} - X_{1,2,3})\right].\end{aligned} \tag{117}$$

For closed shapes (*Cladonema* and *Hydractinia*), we use Eqs. 111-113 to compute the contribution of the basal nematic tension, thickness profile and aboral rigidity modules to both the elongation and the shape polarity (Fig. S10F,G). For species with an oral opening, we used Eqs. 114-117 to compute the contribution of the basal nematic tension, oral geometry, thickness profile and aboral rigidity modules to both the elongation and the shape polarity (Fig. S10A-E).

**Representation of the relative contributions in a ternary diagram:** In Fig. 5K, we compare the relative contribution of the oral geometry, thickness profile and increased aboral rigidity modules to the shape polarity in *Nematostella*, JNK-treated *Nematostella*, *Aiptasia* and *Galaxea*. For that, we define the relative contribution of each module

$$\lambda_m = \frac{|\delta X_m|}{|\delta X_1| + |\delta X_2| + |\delta X_3|}, \tag{118}$$

where labels 1,2,3 refer respectively to the oral geometry, the thickness profile and the aboral rigidity modules. Here, $m = 1, 2$ or 3, such that $\lambda_1 + \lambda_2 + \lambda_3 = 1$. Simulated shapes are then positioned in a ternary diagram, represented by a point $\mathbf{x}$ wtih position:

$$\mathbf{x} = \lambda_2 \mathbf{e}_1 + \lambda_3 \mathbf{e}_2, \tag{119}$$

with $\mathbf{e}_1 = (0, 1)$ and $\mathbf{e}_2 = (1/2, \sqrt{3}/2)$.
